# The eukaryotic translation initiation factor eIF4E reprogrammes the splicing machinery and drives alternative splicing

**DOI:** 10.1101/2021.12.07.471666

**Authors:** Mehdi Ghram, Gavin Morris, Biljana Culjkovic-Kraljacic, Jean-Clement Mars, Patrick Gendron, Lucy Skrabanek, Maria Victoria Revuelta, Leandro Cerchietti, Monica L Guzman, Katherine LB Borden

## Abstract

Aberrant RNA splicing contributes to the pathogenesis of many malignancies including Acute Myeloid Leukemia (AML). While mutation is the best described mechanism underpinning aberrant splicing, recent studies show that predictions based on mutations alone likely underestimate the extent of this dysregulation. Here, we show that elevation of the eukaryotic translation initiation factor eIF4E reprogrammes splicing of nearly a thousand RNAs in model cell lines. Further, in AML patient specimens which did not harbour known splice factor mutations, ∼4000 transcripts were differentially spliced based on eIF4E levels and this was associated with poor prognosis. Inhibition of eIF4E in cell lines reverted the eIF4E-dependent splice events examined. Splicing targets of eIF4E act in biological processes consistent with its role in malignancy. This altered splicing program likely arose from eIF4E-dependent increases in the production of spliceosome components including SF3B1 and U2AF1 which are frequently mutated in AML. Notably, eIF4E did not drive mutation of these factors, only their production. eIF4E also physically associated with many splice factors including SF3B1, U2AF1, and UsnRNAs. Interestingly, eIF4E interacted with both pre-mRNA splicing substrates as well as the resulting product RNAs suggesting that eIF4E could chaperone RNAs throughout splicing as well as other aspects of post-splicing transcript regulation. Importantly, many eIF4E-dependent splice events differed from those arising from splice factor mutation and were more extensive highlighting that these splicing profiles arise from distinct, but potentially overlapping, mechanisms. In all, our studies provide a paradigm for how dysregulation of a single factor, eIF4E, can reprogramme splicing.

## Introduction

Studies into the genomics, epigenetics and transcriptomes of cancer have yielded important insights into its pathogenesis. However, proteomic studies revealed that the transcriptome does not always predict the proteome^1^. This disconnect is due, in part, to post-transcriptional regulation e.g. splicing, nuclear RNA export, and translation. Dysregulation of these events can elevate the production and/or alter the structure and function of proteins involved in all facets of malignancy. Altered splicing is well known to produce a variety of biological impacts on the cells. Splicing is the removal of introns and joining of flanking exons in pre-messenger RNAs (pre-mRNA) and some non-coding RNAs^2^. Most of the splicing is catalyzed by the major spliceosome, an intricate assembly of >150 proteins and 5 uridine-rich small nuclear UsnRNAs (U1, U2, U4, U5 and U6 snRNAs)^2^. This machine recognizes elements in the 5’ splice-site (5’SS), 3’ SS and the branch site to catalyze excision of targeted introns. Alternative splicing (AS) generates greater diversity in the proteome by producing multiple mRNAs from the same pre-mRNA^3^. About 95% of multi-exonic genes undergo AS^4^. AS events include altered selection of the 5’SS or 3’SS, skipped exons (SE), inclusion of mutually exclusive exons (MXE) or intron retention (IR)^4^. AS products can have opposing functions to their constitutive counterparts, lead to transcript and protein mis-localization, generate highly volatile transcripts that are rapidly degraded causing protein loss, or other effects ^3, 5^.

Dysregulation of splicing contributes to hematologic malignancies, solid tumours and genetic diseases^3, 6, 7^. In AML, ∼30% of expressed genes are aberrantly spliced compared to CD34+ cells from healthy individuals^8, 9^. The best characterized modality for driving aberrant splicing in AML involves mutations of splice factors (SF). The most frequently mutated SFs in heme malignancies are SF3B1, SRSF2, and U2AF1 in myelodysplastic syndromes (MDS), and these mutations are associated with progression to AML, with ∼5-10% of AML patients harbouring such mutations^3, 6, 7,10^. Intriguingly, these mutations do not disrupt splicing of all transcripts, but rather have targeted effects. For instance, SF3B1 mutations lead to 83 altered splicing events in AML^11^. Furthermore, in AML, aberrant splicing is much more widespread than the frequency of SF mutations suggesting that there are additional means to modify splicing^12^. For example, two components of the spliceosome (PRPF6 and SF3B1) are elevated in secondary AML specimens without SF mutations, relative to healthy volunteers^13^. In a diverse set of solid tumours, SRSF1, SRSF2, and U2AF2 levels are upregulated^3^. In these AML and solid tumours, the underlying mechanisms for SF elevation are not understood. Identification of modalities by which splicing becomes dysregulated is critical since it will enable us to unravel the molecular basis for reprogramming splicing in the absence of SF mutations which in turn can provide new directions for therapeutic targeting.

Our studies with the eukaryotic translation initiation factor eIF4E suggested the possibility that eIF4E could impact on splicing particularly in AML and other cancers where it is dysregulated. eIF4E is found in the nucleus and cytoplasm where it plays distinct roles in RNA metabolism in both healthy and cancerous cells^14–22^. Interestingly in high-eIF4E AML, eIF4E is not only highly elevated but also enriched in the nuclei at steady-state^14, 23–25^ (Supp Fig1B). This eIF4E phenotype is associated with poor outcomes and indeed can be targeted in AML patients with ribavirin leading to clinical benefit^24, 25^. Indeed, the nuclear distribution of eIF4E is observed prior to ribavirin treatment in high-eIF4E AML patients but then during clinical responses including remissions eIF4E is found mainly in the cytoplasm with its return to the nucleus as an indicator of clinical relapse^24, 25^. Molecular studies revealed that eIF4E entry into the nucleus was mediated by Importin 8^26^. Interestingly, Importin 8 binds the cap-binding site of eIF4E preventing its association with RNAs and thus importing RNA-free eIF4E^21, 26^. Ribavirin, or excess cap analogues, disrupt this eIF4E-Importin 8 interaction preventing nuclear entry of eIF4E and thus provides a molecular basis for the mainly cytoplasmic staining of eIF4E during clinical responses^21, 26^. Upon chemical deactivation of ribavirin, eIF4E can once again bind Importin 8 which allows nuclear entry and underpinning the dysregulated nuclear activities of eIF4E associated with disease progression^21, 26, 27^.

At the molecular level, eIF4E binds the m^7^G cap (referred to as the cap), on the 5’ end of RNAs, an activity conserved between nuclear and cytoplasmic compartments. In this way, eIF4E acts as a cap-chaperone and mediates its activities in the export of selected RNA targets and in the translation of specific transcripts. While these RNAs are capped, other signals referred to as USER codes within the RNAs provide selectivity. For example, in order to be a nuclear RNA export target, RNAs must not only be capped, but contain an element known as an eIF4E sensitivity element (4ESE) within their 3’UTRs. This element recruits the export machinery forming an active export complex with eIF4E-LRPPRC-4ESERNA and CRM1^16, 17, 19, 21^. For translation selectivity, RNAs must contain complex, structured 5’UTRs in addition to their cap^28^. In these cases, eIF4E does not alter the transcript levels of these RNAs. Thus, eIF4E can amplify transcriptional signals for a subset of RNAs with appropriate USER codes by promoting their nuclear export and/or enhancing their translation efficiency^29, 30^.

eIF4E has also been implicated in nuclear RNA maturation including m^7^G capping as well as cleavage and polyadenylation (CPA) of selected RNAs^20, 22, 31^. Thus, eIF4E plays broader roles in mRNA maturation; and this can be viewed as eIF4E acting as an m^7^G cap chaperone^32, 33^. Similar to its selection of nuclear RNA export targets, selectivity arises due to cis-acting elements in the targets RNAs^20, 22^. For the case of both m^7^G capping and CPA, studies revealed a pattern whereby eIF4E: 1. elevates the levels of the CPA and capping machinery through increased export of the transcripts encoding these factors; and 2. eIF4E physically associates with selected components of the capping and CPA machinery^20, 22^. Given these observations, we reasoned that eIF4E similarly impacts splicing of selected RNAs. Here, we demonstrate that eIF4E drives SF production as well as physically interacted with components of the spliceosome. Consistently, eIF4E drives wide-ranging changes to splicing of a broad array of RNAs. In this way, this work uncovers roles for eIF4E beyond amplifying the protein-coding capacity of transcripts, to include molding their physical nature. It also provides a novel paradigm by which to reprogramme splicing. Indeed our studies provide a unique opportunity to dissect mechanisms driving dysregulated splicing independent of SF mutations. In turn, this could substantially expand the use of splicing inhibitors beyond patients harbouring these mutations.

## Results

### eIF4E overexpression impacts protein levels of splice factors through its RNA export function

As a first step to investigate the possible role of eIF4E in splicing, we monitored the impact of eIF4E on levels of SFs derived from the major spliceosome complexes focussing on those associated with cancer e.g. SF3B1 and U2AF1^10^. For this purpose, we initiated our studies in U2OS cells since eIF4E plays well-established roles in capping, CPA, nuclear RNA export and translation here^16–22^. We generated 3 stable 2FLAG-eIF4E or Vector control cell lines as previously descrcibed^22^. Notably, inspection of RNA-sequencing data revealed that neither the parent U2OS cells nor 2FLAG-eIF4E or Vector stable cell lines harbour SF mutations typically associated with cancer e.g. in SF3B1, U2AF1 or SRSF2^10^ (<u>GSE158728</u>). Finally, in U2OS cells we observe endogenous eIF4E in the nucleus in bodies and diffusely throughout the nucleoplasm as well as in the cytoplasm consistent with^16, 19^ (Fig1A). Additionally, the extent of eIF4E elevation in these cells was physiologically relevant; 2FLAG-eIF4E U2OS cells had a ∼3-fold elevation in eIF4E protein levels relative to Vector controls (Fig1B). By comparison, AML specimens had up to ∼8-fold elevation of eIF4E protein levels relative to healthy tissues^24, 25^. To assess SF protein levels, we carried out western blot analyses of total cell lysates. SF3B1, U2AF1, U2AF2, PRPF6, PRPF8, PRPF19, PRPF31, SNRNP200 protein levels were elevated in 2FLAG-eIF4E cells relative to Vector controls (Fig1A). These factors were employed since they are markers of the major spliceosome complexes (Fig1D). Cyclin D1 and Mcl1 served as positive controls while β-actin provides a negative control as it is not an eIF4E target^17–20, 22^ (Fig1A). We next examined whether depletion of endogenous eIF4E reduced levels of these SFs using CRISPR-eIF4E or CRISPR-Ctrl U2OScell lines as described^22^. CRISPR-Ctrl cell lines were generated using guide RNAs to *Galaxidae* coral Azami-green transcripts^22^. CRISPR-eIF4E cell lines were heterozygous for eIF4E as its complete deletion was lethal^22^. As expected, CRISPR-eIF4E cells had reduced levels of PRPF6, PRPF8, PRPF19, SF3B1, SNRNP200, and U2AF2 proteins relative to CRISPR-Ctrl cells (Fig1C). These findings indicated that endogenous eIF4E impacted production of the splicing machinery; and thus, eIF4E’s effects on SFs were not limited to circumstances characterized by eIF4E elevation. In all, we demonstrate that SF production can be driven in an eIF4E-dependent manner. SFs targeted by eIF4E are found in each of the major spliceosome complexes (Fig1D), and included SFs that are often mutated in AML and other cancers e.g. SF3B1 and U2AF1 ^10, 12, 34^. Thus, the impact of eIF4E does not appear restricted to any specific spliceosome complex, but rather, is positioned to influence several splicing steps, and/or suggests that eIF4E is bound to the RNA throughout the splicing process. Further, SF3B1 and U2AF1 which are mutated in many cancers with dysregulated splicing, can also be elevated via eIF4E in the absence of mutation.

**Fig 1.**
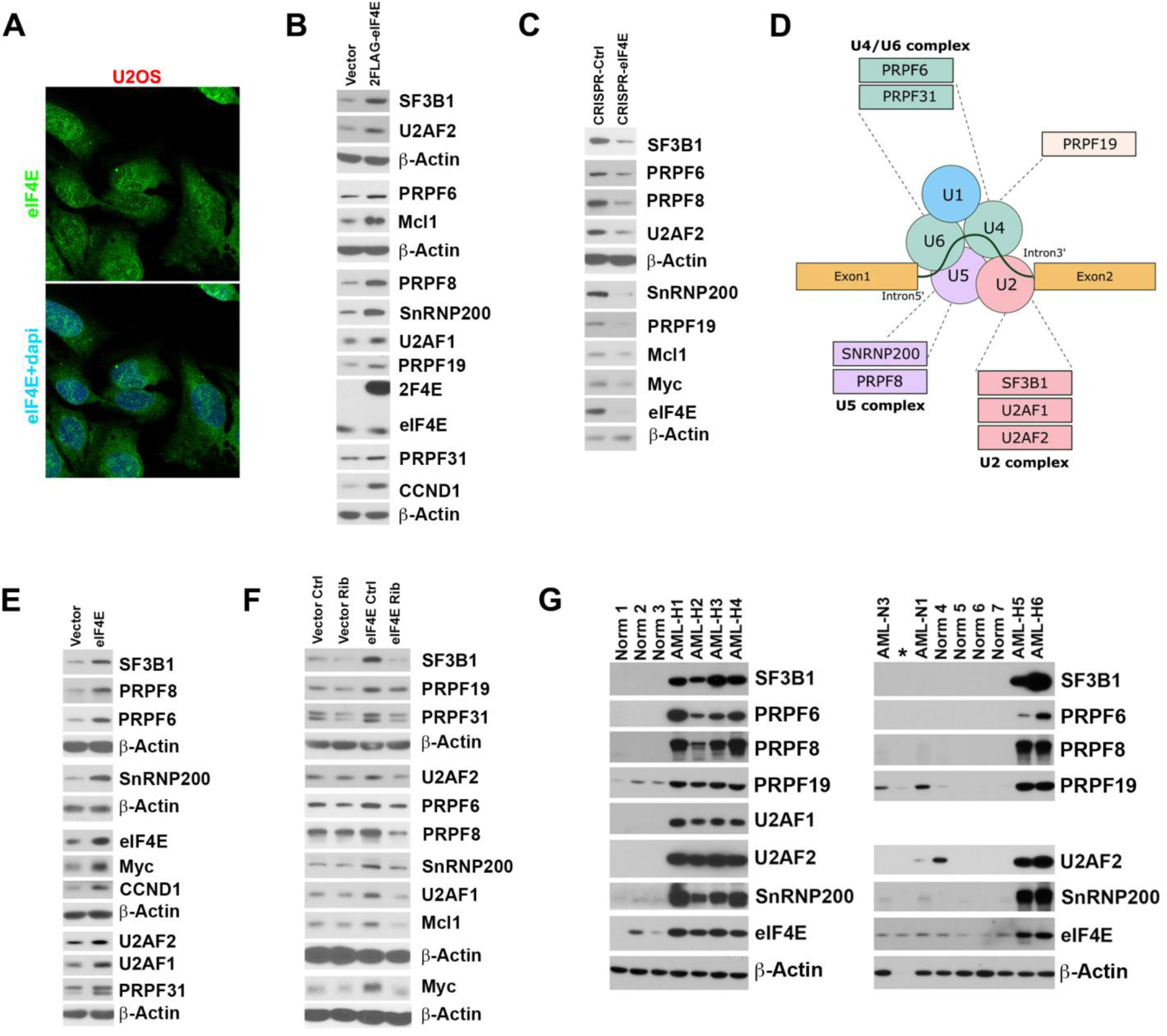
eIF4E regulates expression of the splicing machinery. **A.** Localization of eIF4E in U2OS cells. Confocal micrographs of cells stained with anti-eIF4E antibodies to detect endogenous eIF4E and DAPI as a nuclear marker. Single (eIF4E) and overlaid (eIF4E+DAPI) channels are shown. Micrographs are single sections through the plane of the cells with 63x magnification. **B.** Western blot (WB) analysis of splicing factors for Vector and 2FLAG-eIF4E U2OS cell lines. Myc, Mcl1, and CCND1 served as positive controls, and as a loading control. Both 2FLAG-eIF4E (2F4E) and endogenous eIF4E are shown. Each β-Actin blot corresponds to the above western blots. Experiments were carried out at least three independent times, and one representative experiment is shown. **C.** WB analysis of splicing factors as a function of eIF4E reduction using CRISPR-eIF4E and CRISPR-Ctrl U2OS cell lines. Myc, Mcl1 and CCND1 served as positive controls, and β-Actin was used as a loading control. Each β-Actin blot corresponds to the western blots immediately above. Experiments were carried out at least three independent times, and one representative experiment is shown. **D.** eIF4E is positioned to influence multiple facets of spliceosome activity. Schematic representation summarizing the splicing factors targeted by eIF4E. Proteins are grouped based on their activity and/or association with a specific complex of the major spliceosome. eIF4E also physically interacts with all of the UsnRNAs shown (see Figure 2A and Supp Figure 2). **E.** WB analysis of splicing factors for Vector and eIF4E overexpressing NOMO-1 cell lines. Myc, and CCND1 served as positive controls, and β-Actin as a loading control. Each β-Actin blot corresponds to the above western blots. Experiments were carried out at least three independent times, and one representative experiment is shown. **F.** Western blot analysis of untreated (Ctrl) and Ribavirin treated (Rib) Vector and eIF4E NOMO-1 cell lines. Myc, and Mcl1 served as positive controls, and β-Actin as a loading control. Each β-Actin blot corresponds to the above WBs. Experiments were carried out at least three independent times, and one representative experiment is shown. Ribavirin dose was 10uM which is clinically achievable. **G.** WB analysis of Splicing Factors levels in primary AML samples with high (AML-H) or normal (AML-N) eIF4E levels, as well as bone marrow mononuclear cells from healthy volunteers (Norm for normal). β-Actin was used as a loading control. (* = degraded sample).

We assessed whether SFs were regulated by eIF4E. eIF4E can drive production of proteins through increased cytoplasmic availability of transcripts via increased nuclear export and/or by elevated numbers of ribosomes per transcript thereby increasing translation efficiency. Given the fact that mRNAs coding for factors involved in capping and CPA were eIF4E export targets, we reasoned that eIF4E was positioned to regulate modulate levels of SF proteins by driving export of their corresponding RNAs^20, 22^. First, we carried out eIF4E RNA immunoprecipitations (RIPs) from nuclear lysates since these interactions predict sensitivity to eIF4E-dependent RNA export; by contrast, all examined RNAs bind to eIF4E in the cytoplasm, whether they are translation targets of eIF4E or not and thus this was not prioritized^16–20, 22^. Using nuclear lysates from U2OS cells, we found that endogenous nuclear eIF4E RIPs with *SF3B1, SNRNP200, PRPF6*, *PRPF8*, *PRPF31, U2AF1, U2AF2* (Fig2A) were enriched by ∼2-4 fold versus input. Negative control RNAs *ACTB*, *GAPDH, POLR2A* RNAs and *18*S rRNA were not found in the RIPs while positive controls *MCL1*, and *MYC* were present consistent with previous studies^16, 17, 20, 22^. Given this physical interaction, we examined whether these RNAs were targets of eIF4E-dependent nuclear export by monitoring whether their RNA export was increased in stable 2FLAG-eIF4E cells relative to Vector controls, using the three different cell lines as above. For RNA export assays, cells were fractionated into nuclear and cytoplasmic compartments and RNAs quantified in each fraction using RT-qPCR. As a proof-of-principle, we monitored *SF3B1, U2AF1, U2AF2, SNRNP200, PRPF6, PRPF8* and *PRPF31* all of which had ∼2 fold increased in cytoplasmic/nuclear ratios upon eIF4E overexpression indicating their nuclear export was increased upon eIF4E overexpression (Supp Fig2A). *MYC* and *MCL1* serve as a positive control and were elevated while *GAPDH* and *POLR2A* were negative controls and unchanged as expected (normalized to *ACTB*). Fraction quality was assessed by semi-quantitative PCR monitoring U6 snRNA and tRNA^Met^ for controls for the nuclear and cytoplasmic fractions respectively (Supp Fig2B) ^18^. Typical of eIF4E targets, total levels of the corresponding RNAs were not altered by eIF4E overexpression as observed by RT-qPCR (Supp Fig2C). In all, eIF4E physically interacts with, and promotes the nuclear export of many SF-encoding RNAs providing a mechanism for eIF4E-mediated elevation of these factors.

**Fig 2.**
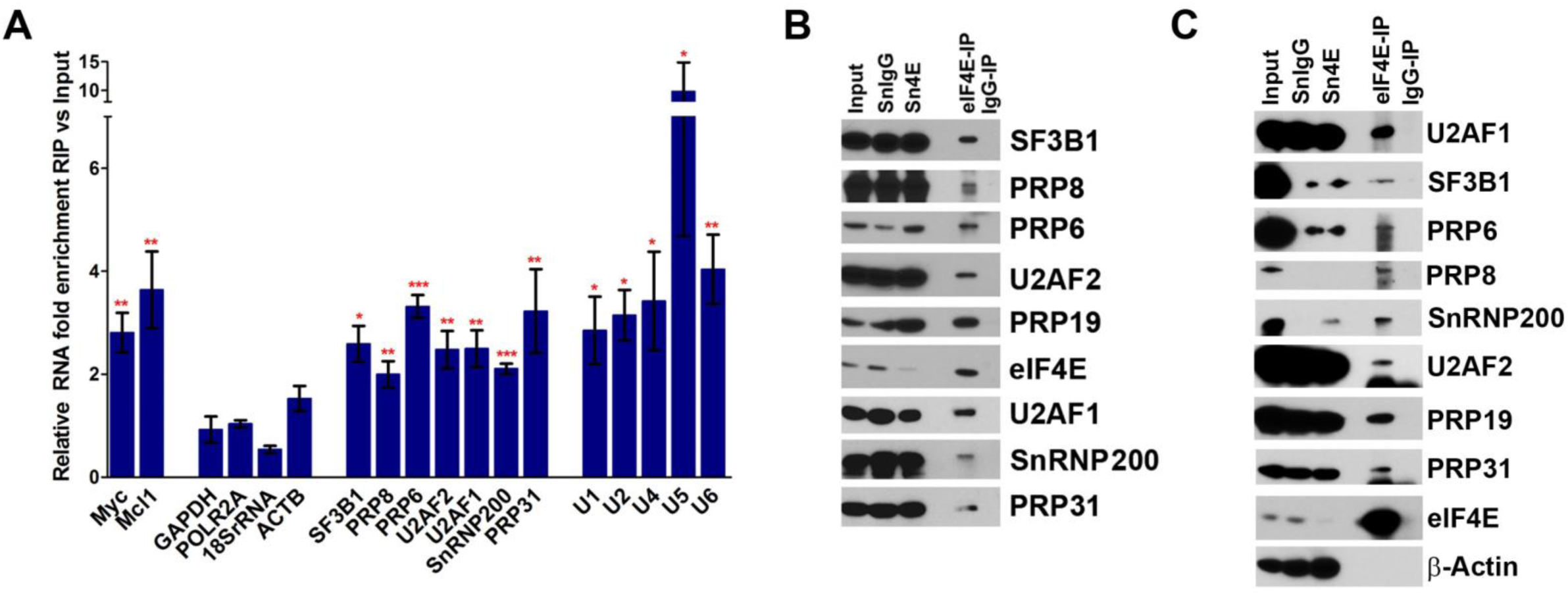
eIF4E regulates expression and interacts with components of the splicing machinery. **A.** The enrichment of mRNAs in RIPs of endogenous eIF4E versus input RNAs from the nuclear fractions of vector control U2OS cells monitored by RT-qPCR. Data were normalized to input samples and presented as a fold change. The mean, standard deviation and p-values were derived from five independent experiments (each carried out in triplicate). Myc, and Mcl1 are established eIF4E nuclear targets and served as positive controls, while ACTB, GAPDH, POLR2A and 18S rRNA were negative controls (* p<0.05, **p<0.01, ***p<0.001). **B.** Endogenous eIF4E co-immunoprecipitated with SFs in the nuclear fractions of U2OS cells. Immunoprecipitations (IP) were carried out using U2OS nuclear lysates and anti-eIF4E antibody (eIF4E-IP) or appropriate IgG control (IgG-IP). IP samples along with Input (2%) and supernatants (Sn) after IPs were analyzed by WB using antibodies as indicated. **C.** eIF4E co-immunoprecipitated with SFs in the nuclear fractions of NOMO-1 cells. IPs were carried out using eIF4E overexpressing NOMO-1 nuclear lysates and anti-eIF4E antibody (eIF4E-IP) or appropriate IgG control (IgG-IP). IP samples were analyzed by WB as above and carried out in three biological replicates.

Given the impact of eIF4E on the production of SFs in U2OS cells, we investigated whether eIF4E drove their production in an High-eIF4E cancer opting to study High-eIF4E AML in which eIF4E has been targeted with ribavirin in early phase clinical trials leading to objective responses including remissions^24, 25^. In these High-eIF4E patients, eIF4E is typically elevated ∼3-8 fold and predominantly localized to the nucleus and characterized by elevated eIF4E nuclear RNA export activity^14, 24, 25^ (e.g. Supp Fig1A). Further supporting this choice of indication, AML is often associated with dysregulated splicing, which is typically attributed to mutation in specific SFs^9, 10, 12, 13, 34^. We used two experimental systems to explore the impact of eIF4E on SFs in AML: the NOMO-1 AML cell line, and primary specimens from AML patients. Given the close link between dysregulated splicing and mutations in *U2AF, SF3B1* or *SRSF2* in AML^9, 10, 12, 34^, we inspected our RNA-Seq data to ensure that NOMO-1 (Supp Fig1C) and the primary AML specimens (GSE67040) examined did not harbour these mutations. In NOMO-1 cells, eIF4E levels are very similar to those in healthy volunteers or Normal-eIF4E AML specimens (Supp Fig1B); thus, these cells provided a unique opportunity to engineer High-eIF4E AML cells, and thus, to genetically dissect the role of eIF4E in an AML context and to compare to our U2OS system to ascertain if this eIF4E mechanisms is conserved in diverse cell types. eIF4E or Vector NOMO-1 cell lines were produced as pools. NOMO-1 eIF4E cells produced eIF4E protein levels and localization similar to High-eIF4E AML specimens (Fig1E, Supp Fig1A bottom panel). We noted that eIF4E overexpression in NOMO-1 cells did not lead to mutations found in AML in the above SFs. We observed elevation of PRPF6, PRPF8, PRP31, SF3B1, SNRNP200, U2AF1, and U2AF2 proteins in eIF4E NOMO-1 relative to Vector NOMO-1 cells (Fig1E and 1F). Further, positive controls Myc, Mcl1, and CCND1 were elevated while the negative control β-actin was unchanged. To assess the impact of eIF4E inhibition, we monitored the impact of the eIF4E-inhibitor ribavirin, which has been used to target eIF4E in early phase clinical trials^24, 25, 35, 36^. We observed reduced SF levels upon eIF4E inhibition with clinically achievable doses of ribavirin in both conditions relative to vehicle treated controls consistent with ribavirin targeting endogenous and overexpressed eIF4E (Fig1F). Thus, overexpression of eIF4E in AML cells led to changes in the splicing machinery similar to those observed in U2OS cell lines suggesting that eIF4E can dysregulate SF production in multiple contexts. Further, its effects on SF production can be reversed by a clinically used eIF4E inhibitor (Fig1F) similar to the reduction observed by CRISPR-4E in U2Os cells (Fig1B).

Next, we examined whether eIF4E status correlated with levels of SFs in primary AML specimens. We isolated de-identified AML patient blasts using FACS^24, 25, 37, 38^ and compared these with bone marrow mononuclear cells or CD34+ cells from healthy volunteers. Analysis of extracted proteins revealed that High-eIF4E AML specimens had elevated levels of PRPF6, PRPF8, PRPF19, SF3B1, U2AF1, U2AF2 and SNRNP200 relative to healthy volunteers or Normal-eIF4E AMLs (Fig1G). Longer exposures of the western blots revealed the presence these factors in the healthy volunteer specimens and normal-eIF4E AML patient specimens (data not shown). Indeed, this highlights the extreme elevation of these SFs in high-eIF4E AML. Thus, elevation of these SFs was not a general feature of AML patient specimens but correlated with eIF4E levels consistent with our studies in U2OS and NOMO-1 cells. While eIF4E is correlated with elevated levels of these factors, it is likely that there are other mechanisms that can drive elevation of SFs, this would be an interesting avenue of future study.

### eIF4E physically interacts with the splicing machinery

Previously, studies showed that eIF4E interacted with RNA processing machinery involved in capping, CPA, export and translation^17, 19–22, 39^. Thus, we examined whether eIF4E similarly interacted with SFs; providing an additional means by which eIF4E could impact splicing. In the nuclear eIF4E RIPs described above, we observed that endogenous eIF4E immunoprecipitated with U1, U2, U4, U5 and U6 snRNAs relative to inputs with a ∼3-10-fold enrichment in nuclear lysates from U2OS cells (Fig2A), as well as in NOMO-1 cells (Fig2D). Endogenous eIF4E did not associate with negative controls (e.g. *GAPDH, ACTB etc*) indicating these RIPs were specific (Fig2A and Supp Fig2D). Additionally, levels of the UsnRNAs were not altered upon eIF4E overexpression in U2OS suggesting that eIF4E did not lead to generation of more spliceosomes (Supp Fig2E). Given UsnRNAs play both structural and catalytic roles in the spliceosome^2^, we investigated whether eIF4E physically associated with protein components of the spliceosome. We found that eIF4E also immunoprecipitated with protein components of the major spliceosome complexes including PRPF6, PRPF8, PRPF19, PRPF31, SF3B1, U2AF1, U2AF2 and SNRNP200 proteins in U2OS (Fig2B) and NOMO-1cells (Fig2C). Notably, these factors were also elevated upon eIF4E overexpression (Fig1) and their corresponding RNAs for those examined bound to eIF4E in nuclear RIPs (Fig2A and Supp Fig2D). In all, eIF4E binds to several SFs and UsnRNAs of the major spliceosome, and elevated SF protein levels in both AML and U2OS cell contexts. Thus, eIF4E is positioned to impact on splicing in different cellular contexts, suggesting this could be a broadly applicable property of eIF4E.

### eIF4E overexpression alone is sufficient to alter splicing programmes

Given these findings, we monitiored the impact of eIF4E on splicing. RNAs from total cell lysates were isolated from the three 2FLAG-eIF4E and Vector U2OS cell lines and subjected to RNA-Seq. Differences in splicing profiles were quantified using replicate Multivariate Analysis of Transcript Splicing (rMATS)^40^. rMATS calculates the “inclusion level differences” for splicing events such as exon skipping (ES), inclusion of mutually exclusive exons (MXE), intron retention (IR) and alternative splice site usage (Fig3A). For example, it calculates the extent an exon is included, i.e. not skipped, in Vector versus 2FLAG-eIF4E cells. In this case, positive values indicate an exon is more included in Vector, and negative values that this exon is more included in eIF4E-overexpressing cells. Here, a value of +1 for SE events indicated that the relevant exon is 100% included in Vector cells and 0% in 2FLAG-eIF4E cells. Many of the same events (129; Supp Table 1) were also observed using EBSeq, which maps exons; but in contrast to rMATS, EBSeq does not provide splice site event information. Thus, we focussed on rMATS. Finally, as stated above, we note that eIF4E-overexpression did not induce mutation in *SF3B1*, *U2AF1* or *SF3B1*, as observed by our RNA-Seq data (GSE158728).We note that RNAs were ribo-depleted to remove ribosomal RNA. Ribodepletion has the advantage over polyA selection that RNAs with short polyA tails or with no polyA tails can be captured in contrast to polyA selection strategies. Given eIF4E has impacts on production of CPA factors^20^ and impacts on APA for hundreds of transcripts (our unpublished observation), we deemed this a necessary consideration.

**Fig 3.**
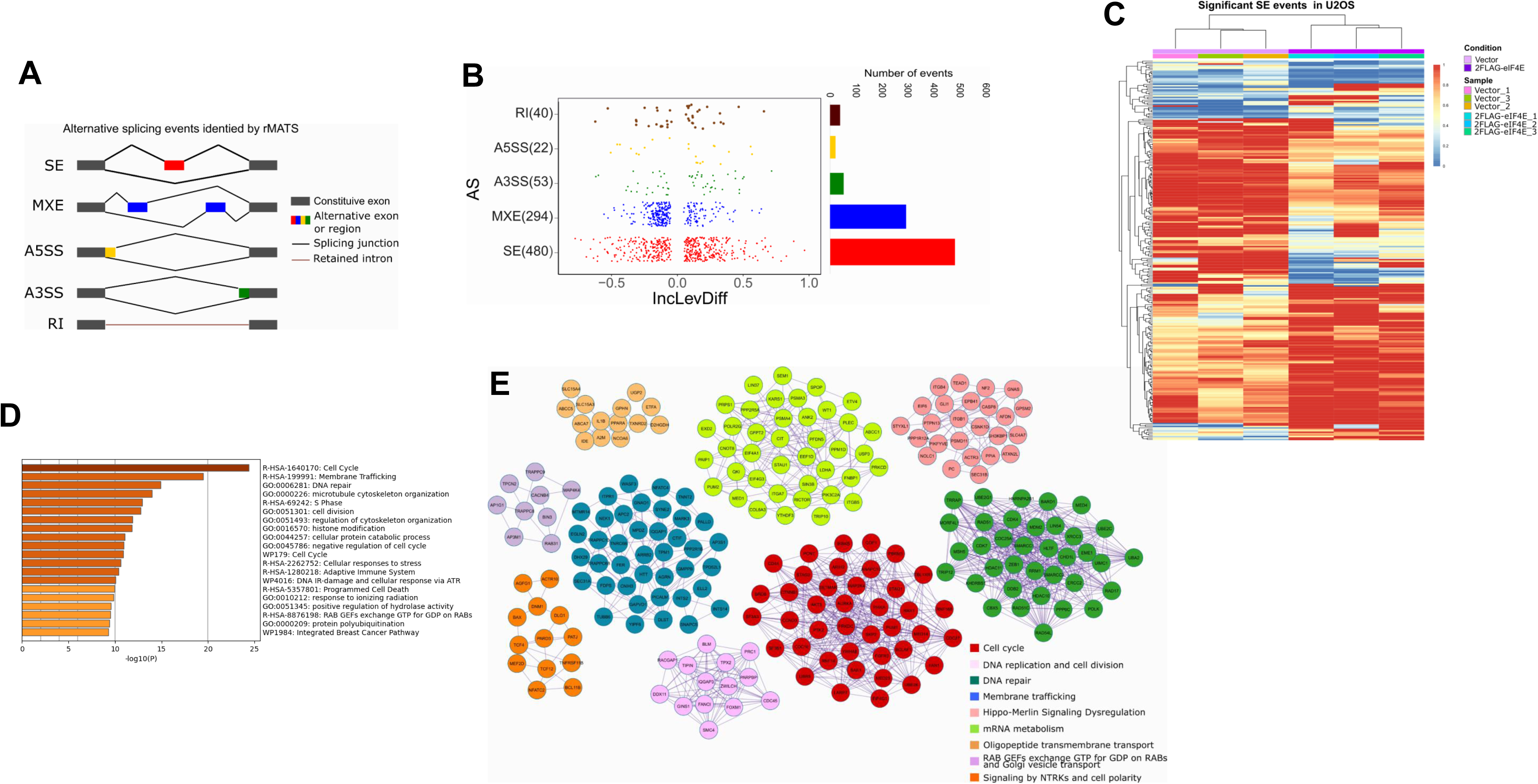
eIF4E overexpression reprogrammes splicing of a subset of transcripts in U2OS cells. **A.** Schematic representation of AS events characterized by rMATS. SE: skipped exon; MXE: mutually exclusive spliced exon; A5SS: alternative 5′ splicing site; A3SS: alternative 3′ splicing site; RI: retained intron. **B.** *Left.* Dot plot showing distribution of Inclusion Level Differences (IncLevDiff) values (IncLev (Vector) – IncLev(2FLAG-eIF4E)) for each splicing category. *Right.* Histogram showing the number of events for each splicing category. **C.** Splicing heatmap showing the values of IncLev for all the Skipping exon (SE) events between Vector (n =3, biological replicates) and 2FLAG-eIF4E (n = 3, biological replicates) U2OS cells. Other events are shown in Supp Fig3A. **D.** Pathway and process enrichment analysis with a bar graph showing top ranked enriched terms for alternatively spliced genes between Vector and 2FLAG-eIF4E U2OS cells. **E.** MCODE networks identified for alternatively spliced genes. Pathway and process enrichment analysis has been applied for each MCODE component and the best-scoring term by p-value is shown.

Using rMATS, we observed that ∼890 splicing events for ∼760 transcripts were altered (FDR-adjusted p-value <0.15; Inclusion differences of >0.05 or <-0.05) or using more stringent cutoffs (p-value <0.1, Inclusion level differences of >0.1 or <-0.1) we observed 555 splicing events for 493 transcripts (Fig3B, Supp Table 2). In the RNA-Seq experiments, a total of 5738 annotated transcripts were detected with >10 TPM indicating that ∼15% of annotated transcripts were differentially spliced in an eIF4E-dependent manner. SE was the most frequent event (∼55%) followed by MXE events (∼30%) (Supp Table 2; Fig3B). More rarely, Intron Retention (IR) and altered 3’/5’ splice sites usage were impacted. The majority of transcripts affected were coding RNAs, but some non-coding RNAs were also impacted (Supp Table 2). Hierarchical clustering analyses based on the inclusion levels across replicates revealed that all types of splicing events (e.g. SE, MXE, IR) segregated solely based on eIF4E levels (Fig3C, Supp Fig3A). Further, the clustering indicated replicates within each group were highly similar. To assess if eIF4E overexpression was associated with repression or promotion of splicing, we analyzed inclusion level differences of the spliced events. Overall, these were roughly equally positive or negative (45% versus 50 % respectively) suggesting a splicing reprogramming of a specific subset of RNAs. In all, these findings support a role for eIF4E as a mediator of alternative splicing (AS) for a subgroup of transcripts; importantly, our studies indicate that eIF4E did not elicit global changes to splicing when overexpressed in U2OS cell lines. Also, eIF4E overexpression did not lead to a global reprogramming of the transcriptome, with only 402 differentially expressed genes in Vector versus 2FLAG-eIF4E U2OS cells^22^. More importantly, only ∼2% of the differentially expressed transcripts (13 targets) undergo AS upon eIF4E overexpression, suggesting that eIF4E-related AS is not generally correlated with RNA levels in these cells (Supp Fig3B, Supp Table 3).

To understand the biological impact of eIF4E-dependent alterations to splicing, we carried out pathway and process enrichment analyses. Enrichment of GO terms for all significant events indicated that eIF4E influenced pathways that could support its oncogenic phenotype (Fig3D, Supp Table 4). Top GO categories include Cell Cycle, Membrane trafficking, DNA repair and Microtubule cytoskeletal organization. Protein-protein interaction enrichment analysis are consistent with the GO terms with the top hits being Cell Cycle, RNA metabolism and Membrane Trafficking (Fig3E, Supp Table 5). The enrichment of RNA metabolism particularly interesting, and suggestive of a positive feedback loop resulting from eIF4E dysregulation and is consistent with its known multiple functions.

### eIF4E levels are correlated with differential splicing programmes in AML

We next examined the relevance of eIF4E-dependent splicing to human diseases characterized by dysregulated eIF4E. We focussed on AML given the extensive studies into dysregulated splicing there and because targeting eIF4E with ribavirin in AML provided clinical benefit including remissions in a subset of AML patients in early stage clinical trials^24, 25^. As a first step, we analysed RNA-Seq data from >450 de-identified AML patient specimens and 17 specimens of normal CD34+ cells derived from human cord blood (www.leucegene.ca) (Fig4). Dividing the entire Leucegene cohort into two groups (above and below median RNA expression of eIF4E), we observed a substantial, and significant, reduction in surival for the High-eIF4E group (Supp Fig4A). For further analysis, we pre-screened specimens by inspection of the RNA-Seq data to ensure they did not harbour the major mutations reported in AML *i.e*. those in *U2AF1, SRSF2* or *SF3B1*. We used the normal CD34+ cells to benchmark eIF4E levels in order to categorize AML specimens into High- or Normal-eIF4E groups (Fig4A). For rMATS analysis, we selected 10 AML specimens with the highest eIF4E levels and 11 with normal eIF4E levels i.e. overlapping with or below levels in normal CD34+ cells (Fig4B). Corresponding clinical data revealed that the 10 High-eIF4E AML patients had drastically reduced overall survival (median survival 136 days) when compared to the 11 patients with lowest eIF4E levels in AML (1396 days, Fig4C).

**Fig 4.**
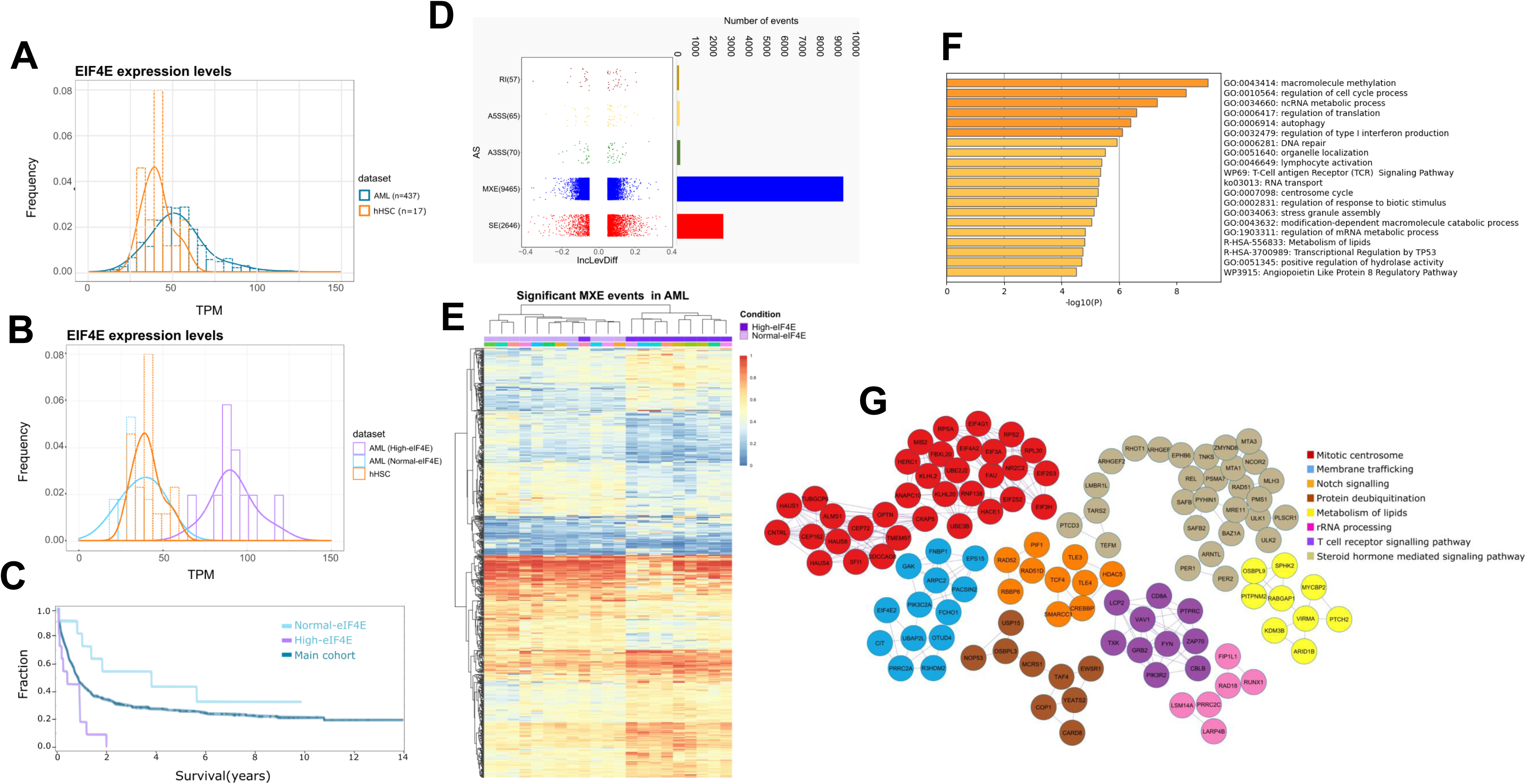
eIF4E levels correlate with a substantial alternative splicing reprogramming in AML. **A.** Graph showing the distribution of eIF4E RNA levels in AML patients’ specimens (blue, AML) in comparison with normal CD34+ cells (orange, hHSC). Data are derived from the Leucegene cohort. **B.** High-eIF4E (purple) and Normal-eIF4E (blue) classification of AML patients’ specimens. eIF4E levels in CD34+ cells from cord blood derived from healthy volunteers is shown for comparison. These specimens were selected both on their eIF4E levels and the absence of SF mutations as observed from the Leucegene RNA-Seq dataset (Supp table 19). **C.** Survival analyses for of High-eIF4E patients and Normal-eIF4E patients for which specimens were studied (Supp table 19). The rest of the cohort is shown for comparison. Log Rank test was performed and shows a significant difference in survival between the High-eIF4E group and the rest of the cohort, as well as between High-eIF4E and Normal-eIF4E groups (p-value < 0.05). **D.** *Left.* Dot plot showing distribution of IncLevDiff values (IncLev (Normal-eIF4E) – IncLev (High-eIF4E)) for each splicing category. *Right.* Histogram showing the number of events for each splicing category. **E.** Splicing heatmap showing the values of IncLev for all the splicing events between Normal-eIF4E and High-eIF4EAML samples. **F.** Pathway and process enrichment analysis with a bar graph showing top ranked enriched terms for alternatively spliced genes between Normal-eIF4E and High-eIF4E AML patients’ samples. **G.** MCODE networks identified for MXE targets between Normal-eIF4Eand High-eIF4E AML patients’ samples. Pathway and process enrichment analysis has been applied for each MCODE component and the best-scoring term by p-value.

We compared the splicing profiles of these High-eIF4E and Normal-eIF4E specimens using rMATS. We observed differences in ∼1600 splicing events impacting ∼1500RNAs (FDR-adjusted p-value <0.1, absolute value of inclusion level differences of >0.1) and if the threshhold is lowered, ∼12000 events for ∼4000 transripts (absolute inclusion value difference of >0.05 and FDR-adjusted p-value<0.15; Fig4D, Supp Table 6).In terms of transcript levels, 8813 genes had an average TPM value >10. Thus alternative splicing of ∼15-50% of detected transcripts corresponds with eIF4E levels, depending on the cutoffs used. Of these events, ∼75% arose due to MXE, ∼20% SE and the remainder were attribuable to retained introns or altered 3’/5’ splice site usage (Fig4D). As for U2OS, High-eIF4E levels correlated with a roughly equal number of promoted and repressed splicing events. The maximum magnitude of the average inclusion level differences across patients was ∼0.4 (corresponding to ∼40% change in population) versus ∼0.8 (∼80%) for U2OS cells (Fig4D versus Fig3B). The reduction in magnitude in primary specimens likely arises due to heterogeneity amongst patients’ transcriptomes^11, 12^. As in U2OS cells, we did not generally observe any correlation between AS reprogramming and changes in RNA levels, since only ∼7% of the alternatively spliced targets were differentially expressed (Supp Fig4C, Supp Table 7, 8).

To establish whether eIF4E levels were the factor most significantly influencing splicing, we conducted unsupervised hierarchal clustering analysis. There was a very tight correlation between eIF4E status in AML patients and MXE events. Specifically, 20/21 AML specimensclustered based on eIF4E levels when monitoring MXE events (Fig4E). Similar analysis of RI events revealed groups clustered on eIF4E levels with the exception of twoHigh-eIF4E patients that clustered with Normal-eIF4E patients (Supp Fig4B). Finally, analyses of SE events indicated two distinct clusters of High-eIF4E AML patients, one cluster was more similar to Normal-eIF4E patients (Supp Fig4B). Interestingly, we note that unlike the overexpression of eIF4E in U2OS cells where only 5 SFs were differentially spliced (Supp Table 2), eIF4E elevation in AML patients correlates with splicing reprogramming of a wide array of spliceosome components (∼20 targets), including *PRPF3, PRPF8, PRPF4B, PRPF39, PRPF31, PRPF6, PRPF40A, SF1, SRSF10, SF3B3, SF3B2, SRSF5* and *U2AF1*(Supp Table 6). Thus, eIF4E appears to have a broader impact on the splicing of SFs in the context of AML. Similar to U2OS cells, we observed that the top GO enrichment terms for eIF4E-dependent MXE targets were highly similar to those in the U2OS datasets: Cell Cycle, DNA repair, Membrane trafficking and RNA metabolism (Supp Fig4F, Supp Table 9). Protein interaction networks was enriched in Mitotic centrosomes, Membrane trafficking and Notch signalling(Fig4G, Supp Table 10).

We reasoned that identification of splicing events conserved across disparate cell types would reveal pan-cancer core networks that underpin, at least in part, eIF4E’s oncogenic effects (Fig3 versus Fig4). A comparison of all eIF4E-dependent splicing targets in U2OS and AML datasets evince a core set of ∼450 common RNA targets (Fig5A, Supp Table 11); absolute value of inclusion level difference>0.05; FDR-adjusted p-value <0.15). Indeed, 60% of targets identifed in the U2OS cells were in common with the AML targets. In terms of transcript expression, AML specimens and U2OS cells had 4705 transcripts in common (>10TPM) and thus about 10% of transcripts that were in common were also eIF4E-dependent splicing targets. Note that core targets were not necessarily characterized by the same splice event in each cell type. GO term analysis of RNA targets revealed that the RNAs enriched between U2OS and AML were a general reflection of those identified in the analysis of either cell-type alone. In all, analyses identifed proteins involved in Cell Cycle, DNA Repair, Membrane Trafficking, Chromatin Modification and RNA metabolism for GO terms (Fig5B, Supp Table 12) and DNA repair, adaptive immune system, MYC activation and others for protein-protein interaction networks (Fig5C, Supp Table 13).

**Fig 5.**
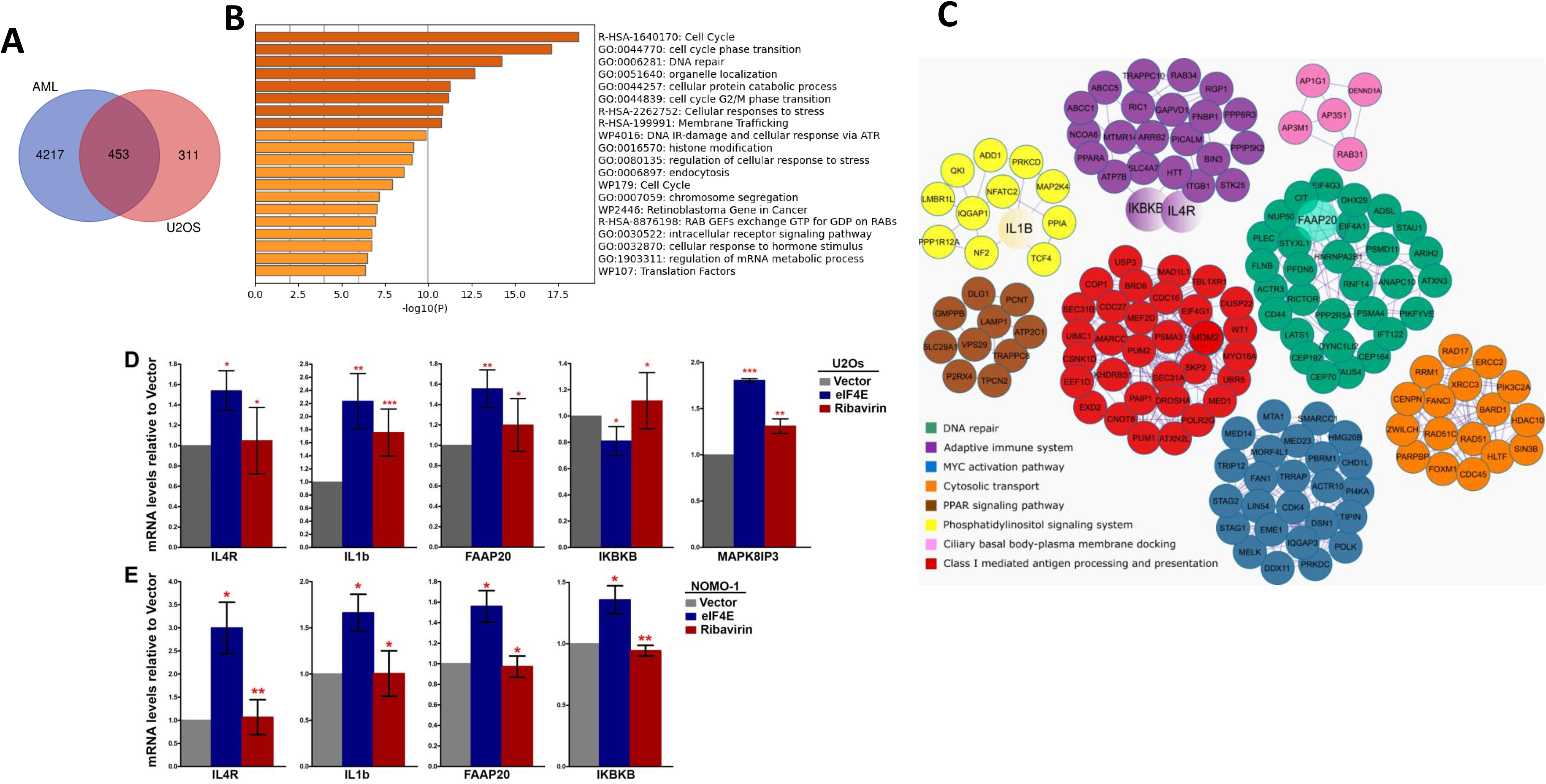
Identification and characterization of a core alternative splicing network targeted by eIF4E in both U2OS and AML contexts. **A.** Venn diagram showing the overlap between genes affected by eIF4E-dependant alternative splicing in High-eIF4E AML patients’ cells and 2FLAG-eIF4E U2OS cells. **B.** Pathway and process enrichment analyses with a bar graph showing top ranked enriched terms for alternatively spliced genes in common between High-eIF4E AML patients’ samples and 2FLAG-eIF4E U2OS cells. **C.** MCODE networks for identified alternative splicing targets common between High-eIF4E AML patients’ samples and 2FLAG-eIF4E U2OS cells. Pathway and process enrichment analysis has been applied for each MCODE component and the best-scoring term by p-value. Common targets selected for subsequent validation by RT-qPCR are highlighted as larger circles (IkBkB, IL4R, FAAP20, IL1B). See main text for full description. **D. and E.** Validation of splicing targets identified by rMATs in U2OS **(D)** and NOMO-1 **(E)** cell lines. RT-qPCR analysis using specific primers for each splicing event normalized to the corresponding total levels of that given transcript (using primers specific to common regions, see supp Table 20). Data were normalized to vector control to calculate fold change. The mean and standard deviation, as well as p-values were derived from three biological replicates (* p<0.05, **p<0.01, ***p<0.001). P-values were calculated between eIF4E overexpressing and vector cells (when the asterisk is over eIF4E), and eIF4E cells treated with Ribavirin vs eIF4E overexpressing untreated cells (for the asterisk over Ribavirin).

### Validation of eIF4E-induced alternative splicing events derived from rMATS analysis using genetic and pharmacological means

Next, we validated several of these core splicing targets in U2OS and for AML using NOMO-1 cells (Fig5D and 5E). We selected events of biological interest to eIF4E’s established oncogenic activities focussing on transcripts from the different protein interaction networks but that were part of the core set of overlapping targets in AML and U2OS cells, and further, prioritized targets with no significant changes to their RNA levels as a function ofeIF4E expression level. Also, we assessed if ribavirin treatment could revert these events to those observed in Vector. Using RT-qPCR, we validated splicing events identified from the rMATS analysis for the following targets as a function of eIF4E overexpression: *IL1B*, *IL4R*, *IkBkB*, *MAPK8IP3* and *FAAP20* in U2OS cells and *IL1B*, *IL4R*, *IkBkB* and *FAAP20* in NOMO-1 cells. As expected, we validated the splicing as seen in rMATS but with one exception. Interestingly, the event that we validated in U2OS cells for *IkBkB* was identified in the AML patients (skipping exon 2) rather than the one predicted by rMATS in the U2OS cells (skipping exon 21) confirming the known limitations of rMATS despite its high efficacy and reproducibility when compared to other event-based AS analysis programs^41^. Next, we examined the impact of eIF4E inhibition with clinically relevant concentrations of ribavirin^25, 26^. We observed in both cell lines that ribavirin treatment reverted the splicing events in eIF4E overexpressing cells *i.e.,* eIF4E-dependent splicing events were reversed back to those observed in Vector cells after treatment (Fig5D and E). To establish the relevance of SFs regulated by eIF4E to these splice events, we employed RNAi methods to knockdown PRPF8, U2AF2 and SF3B1 in eIF4E-FLAG U2OS cells and then compared these to Vector controls. Unfortunately, the single knockdown of any one of these factors led to elevation of the other two factors suggestive of cellular compensation (data not shown). Further, their reduction was associated with massive cell death even under conditions of partial knockdown. Intriguingly, reduction in PRPF8 also decreased eIF4E protein levels. Due to these issues, we were unable to employ this strategy to ascertain which SF components were the most relevant to eIF4E-dependent AS. In all, we validated rMATS predicted eIF4E-dependent AS targets and showed that these splicing effects were reversed by the addition of eIF4E inhibitor ribavirin.

### eIF4E binds not spliced substrate transcripts as well as product RNAs

Given eIF4E impacted splicing of specific RNAs and physically interacted with spliceosome components, we examined whether eIF4E associated with both pre-mRNA splicing substrates and their spliced products. In this way, we investigated whether eIF4E is recruited to the pre-mRNA prior to (or along with the spliceosome) or if these eIF4E-transcript associations were specific to the spliced products. To address this, we carried out nuclear eIF4E RIPs and analyzed RNA content using primers specific to the specific splice event with product-specific primers spanning exon-exon junctions or substrate-specific primers to the relevant intron (or intron-exon boundary) using RT-qPCR. Results were compared to levels of RNAs obtained with primers to both spliced and product RNAs which act as normalization controls. We observe that eIF4E RIPs with transcripts that have not yet undergone the targeted splice event (“not spliced”) as well as those that have undergone that splice event (“spliced”) for *IL1B*, *IL4R*, *IKBKB*, *FAAP20* and *MAPK8IP3* (Fig6A). For example, *IL1B* showed more retained intron in vector cells, and its splicing is enhanced by eIF4E elevation (Fig5D, SuppTable 2). In this case, eIF4E RIPs with both intron-containing *IL1B* and spliced *IL1B* as observed using exon-intron and exon-exon primers, respectively. Thus, eIF4E could play a role in the recruitment of target RNAs. These findings implicate eIF4E in a direct role in splicing for at least some splicing targets. Moreover, these were the first studies to show that eIF4E binds to any pre-mRNAs which has important implications for eIF4E function.

**Fig 6.**
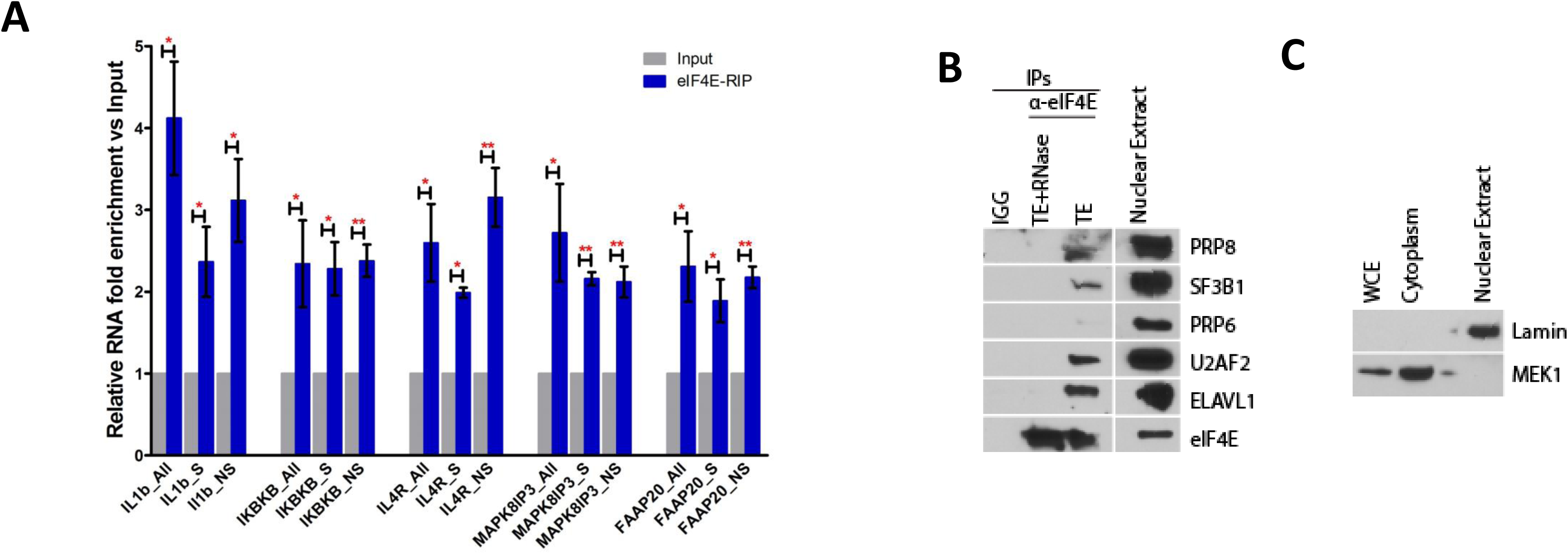
eIF4E binds both splicing substrate and product RNAs, and interacts with the splicing machinery in an RNA-dependant manner. **A.** eIF4E associates with both pre-mRNA splicing substrates and their spliced products. The enrichment of mRNAs for splicing targets in eIF4E-RIPs versus input mRNAs from the nuclear fractions of 2FLAG-eIF4E U2OS cells monitored by RT-qPCR using primers for the specific splice event (S for “spliced”), pre-mRNA specific primers to the relevant intron or intron-exon boundary (NS for “not-spliced”), and primers specific to common regions present in all spliced and not-spliced mRNAs (All). Data were normalized to input samples and presented as a fold change. The mean, standard deviation and p-values were derived from three independent experiments (each carried out in triplicate; * p<0.05, **p<0.01, ***p<0.001). **B.** eIF4E interaction with splicing factors depends on the presence of RNAs. eIF4E immunoprecipitation (IPs) have been performed in U2OS nuclear extract. Rabbit antibodies have been used for IGG and eIF4E. The eIF4E IP has been split in two and treated with TE as control or TE+RNase. After elution, 2% of the nuclear extract and the IPs have been resolved on SDS-PAGE and analyzed by western blotting with the indicated antibodies. Representative of three biological replicates is shown **C.** U2OS cells fractionation for Nuclear eIF4E IPs. The fractionation purity has been tested with indicated antibodies. WCE (whole cell extract).

### The interaction of eIF4E with the splicing machinery is RNA-dependent

We examined whether eIF4E’s interaction with SFs were RNA-dependent to ascertain whether eIF4E’s interaction is mediated via proteins or the substrate RNA. With this is mind, we carried out eIF4E RIPs followed by RNAse treatment of the IPs. RNAse A and T1 were used in combination to efficiently degrade both duplex and loop structures respectively^42^. Endogenous eIF4E was immunoprecipitated from nuclear lysates of U2OS cells and the ability of SFs to bind to eIF4E assessed by western blot (Fig6B). We observed that components SF3B1, U2AF1, U2AF2, PRPF8, SNRNP200 were all present in untreated controls and this interaction was substantially reduced by RNAse treatment. There were no SFs found in the IgG negative controls (Fig6B). Fractionation controls Lamin and MEK1 demonstrate the quality of the nuclear fractions used (Fig6C). Thus, eIF4E require RNAs to interact with all the SFs examined. This suggests that the interaction of eIF4E with the SFs is mediated by the substrate RNA.

### Characterization of eIF4E-dependent AS targets

Above we characterized salient features of eIF4E-dependent AS to understand the basis for selectivity. A key aspect to our understanding of this process is to elucidate the principles of eIF4E-dependent target RNA selection. As a first step, we examined features related to the basic structure of target RNAs with regard to the splice events identified in the rMATS analysis. We monitored exon and intron length, GC content of introns, exon position within the RNA and splice site usage, each as a function of splicing event as well as of positive or negative inclusion level differences. We selected the strongest splice targets to increase the likelihood of identifying commonalities and thus, interrogated both AML and U2OS datasets using the threshold of >0.1 for absolute value of inclusion level differences and FDR-adjusted p-value <0.1. We observed that longer introns were associated with eIF4E-dependent MXE and SE splicing events (Supp Fig5C, Supp Fig6C). Introns were up to ∼9000-15000 bp in average length for eIF4E-dependent events in U2OS cells and ∼6000-9500 bp in AML specimens relative to average intron length for all introns which was ∼4000-5000 bp (Supp Table 14). We observed a trend towards higher GC content for High-eIF4E AML specimens for A3SS, A5SS, and SE events (Supp Fig6E) and for A5SS and MXE events in 2FLAG-eIF4E relative to Vector controls (Supp Fig5E). We did not observe significant alterations in splice site usage in either cell context (Supp Fig5D, Supp Fig6D); consistent with the low number of these events identified by rMATS (Supp Table 2, Supp Table 6). We noted no significant eIF4E-dependent differences for exon length or position (Supp Fig5A, Supp Fig5B, Supp Fig6A, Supp Fig6B). In all, exons involved in eIF4E-dependent splicing events tend to be bracketed by longer than average introns which likely contain relevant sequence features or secondary structure elements that act as USER codes for eIF4E-dependent splicing.

### PRP8, U2AF1 and ARE-binding proteins are identified potential regulators of eIF4E-dependent AS target RNAs

To further dissect sequence elements that impart RNA sensitivity and to determine whether these could be contained within the large introns we identified above, we employed the AURA2 database regulatory enrichment tool to identify RBP-binding sites within eIF4E AS-targets focussing on the core transcripts shared between U2OS and AML groups. We searched for known motifs of RNA-binding proteins and other regulatory cis-elements within 3’ or 5’ UTR regions in the eIF4E-AS core target RNAs. Strikingly, about 60% of the core AS target RNAs contained binding sites for ELAVL1, AGO, eIF4A3, TIA, and IGF2BP which play roles in alternative splicing (Supp Table 15). Also, binding sites for PRPF8 and U2AF2 factors, identified here as both bound to eIF4E and regulated by eIF4E, were found in >20% of core eIF4E-dependent AS target transcripts. Indeed, 60 RNA-binding proteins (RBPs) were predicted to co-regulate >20% of core eIF4E-AS target RNAs and the top-scoring terms associated with these RBPs involved in RNA splicing and processing of capped, intron-containing pre-mRNAs (Fig7A). RNAs contained interaction motifs for several factors indicating that transcripts could be regulated by multiple factors simultaneously. This suggests that a plethora of co-factors could exist to act in eIF4E-AS and that these could be regulated combinatorially as anticipated from the RNA regulon model.

**Fig 7.**
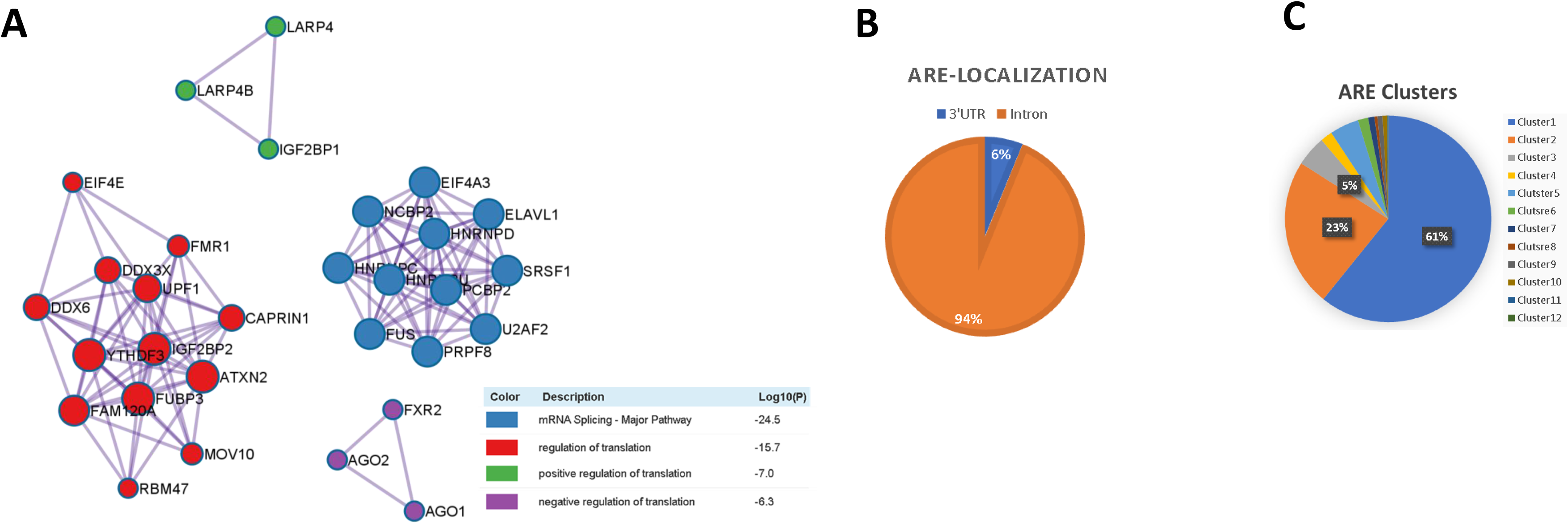
Prediction of AS-eIF4E co-regulatory network and potential AS-eIF4E USER code. **A.** .MCODE networks identified from AURA analysis with 20% coverage of eIF4E-AS targets. Pathway and process enrichment analysis has been applied for each MCODE component and the best-scoring term by p-value is shown. **B.** Pie chart representing the proportion of ARE in eIF4E-AS targets based on their localization. Orange for intronic ARE and blue for ARE in 3’UTR. **C.** Pie chart showing the percentage of each ARE cluster type within eIF4E-AS targets.

Remarkably, many of the RBPs that target >60% of AS-eIF4E target transcripts recognize AU- or U-rich elements including ELAVL1 (also known as HuR), TIA1, TIAL1, AGO (Supp Table 15). These factors were also implicated in alternative splicing and play oncogenic roles consistent with an interplay with eIF4E^43–48^. Of these, ELAVL1 is the best-characterized ARE-binding factor^49–51^ and furthermore, ELAVL1 regulates the RNA stability of eIF4E^52^ suggesting a feedback pathway. Thus, we examined whether endogenous ELAVL1 physically interacted with endogenous eIF4E in the nucleus as a proof-of-principle for interactions with ARE-binding proteins. Indeed, eIF4E immunoprecipitations revealed interactions between eIF4E and ELAVL1 in nuclear fractions but not with IgG controls (Fig6B). Furthermore, the eIF4E-ELAVL1 interaction was sensitive to RNAse treatment, similar to SFs, suggesting that this interaction could be mediated by the RNA splicing target consistent with our above studies (Fig6B).

To ascertain whether the ARE regions were found within the introns, we examined the enrichment of these sequences within eIF4E-dependent AS transcripts as well as determined the location of these elements within the transcripts using the ARE-containing mRNA (ARED) database (Supp Table 16). The ARED analysis demonstrated that ∼80 % of the core AS-eIF4E targets RNAs contain AU-rich elements (Supp Table 16). Cluster 1 and 2 type of AREs constitute ∼60% of elements in eIF4E-AS targets. About 94% of identified AREs were found in introns^49^. Inspection of targeted introns of our validated targets*, IL4R, MAPK8IP3, FAAP20* and *IKBKB* revealed these contained AREs which is consistent with interactions with ELAVL1 (Supp Table 16). In all, our studies suggest that the presence of AU rich elements within introns contribute to selectivity of eIF4E-AS target RNAs. This will be further dissected in future.

## Discussion

In this report, we unearthed a novel paradigm for reprogramming splicing which has far-reaching impacts *e.g.,* ∼15% of transcripts in U2OS cells and ∼15-50% of transcripts in AML patient specimens. Underpinning these changes, eIF4E drives wide-scale, simultaneous alterations to the production of splicing factors, and at the same time physically interacts with the splicing machinery including the UsnRNAs. eIF4E-dependent alterations do not induce well known mutations in SFs, alter UsnRNA levels or generally alter target transcript levels. Thus, classical genomic and transcriptomic approaches would be blind to eIF4E’s impacts on splicing and likely explain why these impacts have not been previously observed.

In terms of mechanisms, we observed two non-mutually exclusive means by which eIF4E is positioned to influence splicing (Supp Fig8): through physical interactions with splicing components and substrate RNAs, and/or by regulation of SFs. In terms of the former mechanism which we denote the direct model, eIF4E is positioned to influence splicing through RNA-mediated physical interactions with snRNPs and splicing substrate pre-mRNAs. In this model, eIF4E binds the m^7^G cap of AS target intron-containing RNAs and stays associated with the RNA through the splicing steps (Supp Fig8). Consistent with this model are the observations that eIF4E associates with SFs in a RNAse-sensitive manner (Fig6B) and that eIF4E binds both the substrate pre-mRNA and product mRNAs (Fig6A)). After splicing, eIF4E is positioned to escort the newly spliced capped-RNAs to other processing steps, to the export machinery and/or possibly is exchanged for other nuclear cap-binding proteins such as the nuclear cap-binding complex (CBC). In this way, eIF4E acts as a cap-chaperone^32, 33^ (Supp Fig8), in much the same manner that the CBC does for bulk pre-mRNA during splicing^53–55^. Interestingly, CBC is linked to splicing of the first intron of target transcripts^54^, a preference we did not observe for eIF4E-sensitive transcripts (Supp Fig5 and 6).

The direct model gives rise to considerations of how RNAs are selected to undergo eIF4E-dependent AS rather than CBC-dependent, bulk pre-mRNA splicing (Supp Fig8). Indeed, the CBC and eIF4E compete for capped RNAs in the nucleus^19, 33^ and we observe a CBP20-eIF4E interaction suggesting that capped RNAs could be exchanged between these factors (Mars et al, unpublished observations). The best-described example of this cap competition comes from studies of eIF4E-dependent nuclear RNA export^19^. In this example, cellular RNAs containing specific USER codes*, e.g*. 50-nucleotide 4ESE element in their 3’UTR, are enriched in nuclear eIF4E RIPs and favour the eIF4E-dependent export pathway in high-eIF4E conditions; while in normal eIF4E conditions, these transcripts are more equally distributed between eIF4E and CBC immunoprecipitations and resultant export pathways used^16, 19^.

We propose a similar model for selection of RNAs for eIF4E-dependent splicing (Supp Fig8). eIF4E-dependent AS is likely favoured by RNAs which contain USER codes in their long introns which subsequently enlist factors like ELAVL1 and/or SFs. Indeed, our discovery that 94% of eIF4E-dependent AS targets are enriched for ARE in their introns provides a strong basis for this hypothesis as does the RNA-dependent interaction between eIF4E and the ARE-binding protein ELAVL1 (Fig6B). It is necessary to further characterize splicing USER codes that can recruit or exclude appropriate factors permitting increased (or reduced) eIF4E-dependent AS. Further defining these elements is a major undertaking that is an important future direction of this work. In all, our data support a model whereby eIF4E acts as a cap-chaperone and USER codes within the target RNAs recruit factors that favour eIF4E-dependent AS over CBC-dependent, bulk splicing. Additional experimentation is required to further support this model. Indeed, it is noteworthy despite many years of study the exact role that CBC plays in splicing is still not fully understood, even though its requirement for many splicing events is clear^53, 54, 56^.

Another consideration relevant to the direct model is that splicing is not carried out by a single complex but rather by a series of snRNPs which do not simultaneously bind a given splice site^2^. For example, U1 and U2 are recruited to define the splice site initially, while during the course of spliceosome remodelling U1 and U4 are generally removed while U2, U5 and U6 snRNPs comprise the catalytic complex^2^. Yet, we observed that eIF4E binds to representative factors of all 5 major snRNPs including SFs as well as their cognate UsnRNA factors (Fig2). The interaction of multiple splicing complexes with eIF4E could arise due to two non-mutually exclusive reasons: 1. eIF4E-RNA complexes represent a snapshot of a population of the same species of transcript at different stages of splicing and/or 2. eIF4E-RNA complexes are occupied by snRNPs at the eIF4E-dependent targeted AS splice site, but U1 and U2 snRNPs subsequently mark splice sites on unrelated intron-exon boundaries for other splicing events. Indeed, other groups have observed the recruitment of U1 and U2 after the spliceosome assembles on other sites to presumably enhance splicing assembly for upcoming splice events^57^. These possibilities are difficult to disentangle experimentally and likely to co-exist. Another important consideration is our observation that eIF4E RIPs with intron-containing target RNAs (Fig6A). Indeed, typically splicing occurs on the second timescale^58^. Thus, the stability of these eIF4E-intron-containing RNA complexes appearing at steady state, suggest the complex is stable beyond the time it takes for splicing to occur. Another possibility which is not mutually exclusive is that the splicing of longer introns is slower leaving eIF4E on the RNAs for a longer time or that new not-spliced transcripts are constantly added after each splicing event. Thus, eIF4E’s interaction with the intron-containing RNA may imply that eIF4E can sequester these RNAs at specific sites in the nucleus until point when these target RNAs undergo splicing. Along these lines, it is not yet known whether eIF4E participates in co-transcriptional and/or post-transcriptional splicing. Post-transcriptional splicing occurs at splicing speckles and is characterized by longer introns and slow splicing^59^. eIF4E appears to favour splicing changes to RNAs with longer introns (Supp Table 14) implicating eIF4E in post-transcriptional splicing perhaps as nuclear speckles. Future studies will be needed to dissect these possibilities.

In addition to its direct role, eIF4E can simultaneously influence splicing indirectly through its impact on production of SFs. We refer to this as the indirect model. The capacity of eIF4E to modulate the levels of some SFs suggest that it modifies the composition and subsequent activity of the splicing machinery, potentially producing specialized spliceosomes to favour certain splicing outcomes and to repress others. Interestingly, SFs such as SF3B1 and U2AF1, which are eIF4E direct and indirect targets (Fig1, Fig2), are present in sub-stoichiometric quantities relative to the enzymatic machinery of the spliceosome^2^. This positions eIF4E to influence production of spliceosomes with different RNA preferences. Notably, eIF4E elevation leads to enhancement or reduction depending on the splicing event, and thus, is not a simple product of the cells having elevated levels of the splicing machinery. Indeed, eIF4E did not elevate levels of UsnRNAs themselves, suggesting that eIF4E does not increase the number of spliceosomes but rather impacts their composition to modify their targeting preferences. Finally, we observe substantial overlap between the direct and indirect models of eIF4E mediated AS. For instance, eIF4E both increases the levels of, and binds to, SFs from all the major snRNPs (Fig1, Fig2). Indeed, it is not possible to disentangle the direct and indirect models experimentally. However, this mechanistic entanglement suggests strong eIF4E-dependent biological feedback driving the substantive AS changes we observed. Indeed, taken together with the mechanistic similarity for eIF4E-dependent capping and CPA^20, 22, 31^, eIF4E is positioned to broadly impact all the major aspects of RNA maturation through this pattern of effects.

We note that eIF4E-dependent AS specificity is not based *a priori* on sensitivity to eIF4E regulation at other levels of RNA processing. Thus, we posit that there are cis-acting elements in the RNA targets that sensitize transcripts specifically to eIF4E-dependent splicing and not universally to all eIF4E-related processes. For instance, *CCND1, MCL1* and *MYC* are found in eIF4E nuclear RIPs and are targets of eIF4E-dependent capping and nuclear export but were not observed as eIF4E-dependent splicing targets according to the rMATS analysis (Supp Table 2, Supp Table 6)^16, 17, 22, 60^. A comparison of datasets reveals that six targets of eIF4E-dependent capping^22^ are also capping targets (*MDM2*, *AGRN*, *TUBB6*, *CATSPER1*, *CTNNB1* and *ERCC*2; Supp Fig7B, Supp Table 19) indicating that co-regulation of capping and splicing can occur.

Previous studies suggested that mutations in *SF3B1*, *U2AF1* and *SRSF2* do not occur simultaneously in the same AML cell likely because they are synthetically lethal^10^. Strikingly, we observed that eIF4E is positioned to simultaneously dysregulate these factors through their elevation and/or through their interactions with eIF4E. Interestingly factors such as U2AF1 are considered to be substochiometric^2^, and thus eIF4E’s elevation of these factors can result in reprogramming of splicing without global alterations to the machine. Given the pervasive effects of eIF4E, it is extremely unlikely that its splicing signature would be attributable to elevation of and/or interaction with any single SF. Consistent with this notion, in secondary AML (sAML) specimens without SF mutations, PRPF6 and SF3B1 protein levels were elevated relative to healthy volunteer specimens^13^. Only 363/4670 of eIF4E targets (363/963 for secondary AML targets) were in common between High-eIF4E AML and sAML specimens (Supp Fig7C, Supp Table 17). Thus, the elevation of PRPF6 and SF3B1 alone was not sufficient to recapitulate the much broader ranging eIF4E-dependent splicing profile. Moreover, splicing patterns arising from eIF4E overexpression were not simply a recapitulation of those observed in AML cells with SF mutations. For instance, a comparison of splicing events arising in High-eIF4E AML cells relative to AML cells with *SF3B1* hotspot mutation(SF3B1^K700^ and other SF3B1 mutations were also included)^11^ showed only a modest overlap (47/4670) of eIF4E targets but included a substantial fraction of SF3B1 targets (47/83 of SF3B1) (Supp Fig7C, Supp Table 20). Recent findings indicate that Myc hyperactivation stimulates splicing factor SF3A3 translation through eIF machinery, leading to a metabolic reprogramming that amplifies Myc’s oncogenic potential^61^. A comparison of splicing targets arising due to Myc dysregulation with eIF4E targets reveal a common 146/764 of targets for eIF4E and 146/1442 of Myc targets (Supp Fig7D, Supp Table 18). In all, these analyses suggest that eIF4E dysregulation of splicing is not readily recapitulated by any single downstream target of eIF4E. These studies with eIF4E provide a possible mechanism for simultaneous dysregulation of multiple splicing factors which has been reported in some solid tumours. For example, PRPF8, SRSF1, SRSF2, and U2AF2 protein levels are upregulated, rather than mutated, in some solid tumours^3^ and thus eIF4E could provide a mechanism for their elevation, a hypothesis that could be tested in future. Consistent with the findings presented here, others showed that modulation of the core components of the splicing machinery such as SF3B1 do not elicit global effects on splicing but rather target specific pre-mRNAs^62^.

In all, the collective alterations to the SF content in combination with eIF4E’s interaction with the splicing macheinry and substrate RNAs likely combinatorially drive the broad-ranging altered splicing patterns observed in U2OS and AML cells. Targeted RNAs act in processes that could underpin, at least in part, the oncogenic activities of eIF4E, and likely represent an eIF4E-dependent splicing RNA regulon. Moreover, we show for the first time that eIF4E binds to pre-mRNA as well as mature transcripts. In summary, our study reveals a novel mechansims for dysregulated splicing and further showcases that eIF4E can both amplify and re-write the RNA message.

## Acknowledgements.

We are grateful for assistance from Genomics Platform at IRIC. Primary human specimens were collected and analyzed by the Banque de cellules leucémiques du Québec (BCLQ), supported by the Cancer Research Network of the Fonds de Recherche du Québec - Santé (FRQS). KLBB acknowledges funding from Leukemia and Lymphoma Society USA and Canada, the National Institutes of Health, the Canadian Insitutes for Health Research and, holds a Canada Research Chair in Molecular Biology of the Cell Nucleus; and MG from The Cole Foundation and Baumgartner fellowships.

## Materials and Methods

### Plasmids, Antibodies and Reagents

pcDNA-2Flag-eIF4E wild-type and MSCV-pgk-GFP-eIF4E constructs were previously described^14, 17, 60^.Antibodies for immunoblotting: Mouse monoclonal anti-eIF4E (BD Biosciences), mouse monoclonal anti-β-actin (Sigma Aldrich), rabbit polyclonal anti-Mcl-I (S-19) (Santa Cruz), mouse monoclonal anti-HSP90α/β (F-8) (Santa Cruz), rabbit polyclonal anti-Myc (ab32072 Abcam), rabbit polyclonal antiCyclinD1 (ab134175 Abcam), rabbit polyclonal SF3B1 (Cell Signaling), rabbit polyclonal anti-PRP8 (Bethyl), rabbit polyclonal anti-PRP6 (Bethyl), rabbit polyclonal anti-SnRNP200 (Bethyl), rabbit polyclonal anti-U2AF1 (Bethyl), rabbit polyclonal anti-PRP31 (Bethyl), mouse monoclonal anti-PRP19 (Santa Cruz) and mouse monoclonal anti-U2AF2 (Santa Cruz).

### Cell Culture

U2OS cells were obtained from ATCC, and maintained at 37°C and 5% CO2 in Dulbecco’s modified Eagle’s medium (DMEM) (ThermoFisher Scientific) supplemented with 10% fetal bovine serum (FBS) (ThermoFisher Scientific) and 1% penicillin-streptomycin (ThermoFisher Scientific). NOMO-1 cells (obtained from DSMZ) were maintained at 37°C and 5% CO2 in Roswell Park Memorial Institute (RPMI) 1640 medium (ThermoFisher Scientific) supplemented with 10% FBS and 1% penicillin-streptomycin. Vector and 2FLAG-eIF4E wildtype U2OS cell lines were generated as described previously^17, 60^, and maintained as U2OS cells with addition of G-418 (1 mg/mL, Wisent Bioproducts). MSCV-pgk-GFP-eIF4E wildtype were used for retroviral transduction of NOMO-1 cells as in (Topisirovic et al 2003)^14^. Internal ribosomal entry site (IRES) is placed between GFP and eIF4E and thus eIF4E is not produced as a fusion protein. Transduced cells were isolated using FACSAria cell sorter (BD Biosciences). Cell lines were authenticated using STR profiling (Wyndham Forensic Group). Cultured cells were routinely checked to ensure that there was no mycoplasma contamination by PCR (Sung et al, 2006)^63^. U2OS cells were treated with 20mM ribavirin (Kemprotec, UK) for 72h, and NOMO-1 cells were treated with 10μM ribavirin for 72h.

### Primary specimens

For all patients in this study, written informed consent was obtained in accordance with the Declaration of Helsinki. All specimens were deidentified. Patient specimens for analysis of splicing factors levels were obtained from the Banque de Cellules Leucémiques du Québec with institutional ethics approval. Samples for WB in Fig1G (left panel) and AML-High in Supplementary Fig1B were samples obtained from AML patients prior to treatment with ribavirin combinations in clinical trial (ClinicalTrials.gov registry is NCT02073838). This study received institutional review board and Health Canada approval. Blasts were isolated using flow cytometry as described^24^. Normal specimens were obtained from ATCC. Samples used for rMATS analysis are listed in Supp Table 21

### Immunofluorescence and Laser-Scanning Confocal Microscopy

U2OS cells were grown on 4 well glass slides (Millicell EZ SLIDE 4 well glass, Millipore Sigma PEZGS0416). After washing three times in 1× PBS (pH 7.4), cells were fixed in pre-chilled Methanol at −200C for 10 min, and air dried at room temperature for 30min. NOMO-1 cells were harvested and washed three times in 1xPBS in eppendorf tubes, with centrifugation of 5minutes at 1200 rpm. Washed pellets were resuspended in 1xPBS at 50000 cells/10μl, and 20μl cell suspensions were spotted on glass slides. After drying for 30 minutes at room temperature, NOMO-1 cells were fixed in pre-chilled Methanol at −200C for 10 min, and air dried at room temperature for 30min. After drying, slides were blocked for 1 h in Blocking solution (10% FBS and 0.1% Tween 20 in PBS), and incubated with eIF4E-FITC (BD Biosciences) directly conjugrated antibody diluted in Blocking solution (1:50) overnight at 4 0C. After washing three times in PBS, cells mounted in antifade mounting medium with DAPI (Vector Laboratories). The cells were washed four times with PBS and mounted in mounting media with DAPI (Vector Laboratories, H-2000). Analysis was carried out using a laser-scanning confocal microscope (LSM700 META; Carl Zeiss, Inc.), exciting 405 nm and 488 nm with 63x oil objective and numerical aperture of 1.4. Channels were detected separately, with no cross talk observed. Confocal micrographs represent single sections through the plane of the cell. Images were obtained from ZEN software (Carl Zeiss, Inc.) and displayed using Adobe Photoshop CS6 (Adobe).

### Cellular Fractionation and RNA export assay

About 5 × 10^7^ U2OS cells were collected and washed twice in ice-cold PBS (300 x g for 3–5 min) and then resuspended with slow pipetting in 0.5 mL of lysis buffer B (10 mM Tris (pH 8.4), 140 mM NaCl, 1.5 mM MgCl2, 0.25% Nonidet P-40, 1 mM DTT, 100 U/mL RNase inhibitors). The lysate was centrifuged at 1000 x g for 3 min at 40C, and supernatant (cytoplasmic fraction) was transferred into a fresh microtube. The pellet (nuclear fraction) was resuspended in 1 Volume of lysis buffer B and transferred to a round-bottomed polypropylene tube, and 1/10 volume of detergent stock (3.3% sodium deoxycholate, 6.6% Tween 40 in DEPC H2O) was added with slow vortexing (to prevent the nuclei from clumping) and incubated on ice for 5 min. The suspension was transferred to a microtube and centrifuged at 1000 x g for 3 min at 40C. Supernatant (post-nuclear fraction) was added to cytoplasmic fraction. RNA was extracted from the different fractions by adding TRIzol reagent (ThermoFisher Scientific) and isolated using Direct-zol RNA Mini-prep Kit (Zymo Research).

### Co-Immunoprecipitation and RIP

Nuclei isolated using the Cellular Fractionation protocol were rinsed 2x with 1xPBS and fixed with 1% PFA for 10min at RT with rotation, quenched 5min with 0.15M Glycine (RT with rotation), then washed 3 times with 1xPBS and lysed in 0.5ml NT-2 buffer by 3 times 6 seconds bursts (with 30 second pause between each burst) using microtip at 25% power (Sonic Dismembrator Model 500, Fisher, Max Output 400W). NT-2 buffer: 150mM NaCl, 50mM Tris-HCl (pH 7.4), 2.5mM MgCl2, 0.05% NonidentP-40, 8 supplemented with 1mM DTT, 1x protease inhibitors without EDTA, 200U/ml RNaseOut. Nuclear lysates were centrifuged at 10 000 x g for 10min, and supernatants were transferred into fresh tubes. After adjusting the concentration to be no more than 1mg/ml, nuclear extracts were pre-cleared with 50 μL protein G conjugated superparamagnetic beads (Dynabeads Protein G, ThermoFisher Scientific) for 30 min at 40C. Pre-cleared lysates (1mg) were incubated with 10μg of anti-eIF4E antibody (RN001P, MBL) or 10 μg of appropriate IgG as a control, and 0.5 mg/ml yeast tRNA (SigmaAldrich), overnight at 40C with rotation. After ON incubation, 50μl of Dynabeads were added and incubated for additional 3h at 40C with rotation. Beads were washed once with NT-2 buffer supplemented with 1mg/mL heparin (Sigma-Aldrich) for 5min at 40C with rotation, and an additional six times with NT-2 buffer with 300mM NaCl. After washing, beads were resuspended in 2xLaemmli Buffer with β-mercaptoethanol and incubated for 5 min at 980C. Co-immunoprecipitated proteins were resolved on SDS-PAGE and visualized by Western blotting. To isolate RNAs from immunoprecipitated reactions, beads were resuspended in Elution Buffer (100 mM Tris-HCl (pH 6.8), 4% (w/v) SDS, 20% (v/v) glycerol, 12% (v/v) β-mercaptoethanol, and incubated for 5 min at 980C. RNA were isolated using TRIzol reagent (ThermoFisher) and Direct-zol RNA Micro-prep Kit (Zymo Research).

### Direct eIF4E IP and RNase Elution

Cells was collected and fractionated as described previously^64^. After fractionation, nuclei have been lysed by sonication (20% power, 3×6s with Sonic Dismembrator Model 500, Fisher, Max Output 400W) in 0.5mL per 4×10 7 cells of NT-2 buffer supplemented with 1x protease inhibitors. Lysates has been cleared by centrifugation at 10000xg 10min at 4°C, transferred into clean tube and the concentration has been determined by BCA assay before adjusted to 1mg/ml. Nuclear Lysate has been cleared with 30ul of Dynabeads G per mg of extract at 4°C for 40min. For 1mg IP, 33ul of Dynabeads G (Invitrogen) have been pre-incubate with 10ug of anti-eIF4E (MBL) or rabbit IGGs at RT for 20 min. After 5 washes, beads are resuspended with pre-cleared lysate and incubated with rotation at 4°C o/n. IPs has been washed 5 times with 1ml of NT-2 buffer. IPs has been re-suspended in 1x beads volume of TE (10mM Tris pH8, 1mM EDTA) +0.5ng/ul RNaseA (Qiagen 158922) and 0.05U/ul of RNaseT1 (Sigma R1003) and incubate at RT for 15min with agitation, supernatant has been kept in a new tube. IPs has been rinsed with 1x beads volume of TE and supernatant have been pooled with the eluate. IPs has been washed once with NT-2 bufferand beads has been re-suspended in 2x LB and incubate at 95°C for 10min. Samples has been resolved by SDS-PAGE and visualized by Western Blot.

### Reverse transcription and quantitative PCR

RNA samples were reversed transcribed using SuperScript VILO cDNA synthesis kit (for RIP experiments) (ThermoFisher Scientific) or MMLV reverse transcription kit and oligo-dT or Random hexamers (ThermoFisher Scientific). QPCR analyses were performed using SensiFastSybr Lo-Rox Mix (Bioline, MA, U.S.A, Cat# BIO-94020) in Applied Biosystems Viia7 or QuantStudio7 thermal cyclers using the relative standard curve method (Applied Biosystems User Bulletin #2). All the primers are listed in Supp 20 and all conditions were described previously^60^.

**Western blot analysis** was performed as described previously^38^. Blots were blocked in 5% milk in TBS–Tween 20. Primary antibodies were diluted in 5% milk.

### Analysis of alternative splicing

Alternative splicing events were detected in RNA-seq data and the probability that the differences in isoform abundance were evaluated statistically with the help of rMATS version 4.1.1. Sequenced fragments were treated as non stranded to allow a uniform treatment of both stranded and non stranded samples and variable read lengths were allowed in the analysis. JC and JCEC quantifications have been combined for the complete analysis. Events with a false discovery rate below 0.1 or 0.15 and an absolute inclusion level higher than 0.1 or 0.05 were considered significant and carried further in the analysis.Maxent was used to evaluate the strength of significant splice site events. MAXENT ranges considered for the analysis were : Strong: MAXENT score >= 7, Intermediate: MAXENT score >= 3 and < 7, Weak: MAXENT score < 3.

### Data analysis and visualization

DESeq2 version 1.30.1^65^ was used to normalize gene readcounts (non stranded) for the 21 AML samples. Significant differentially expressed genes (DEGs) are typically those with padj lower than 0.05 and an absolute fold change > 2. **Data visualization, graphics and plots** were made using R package ggplot2 and related packages digest, glue, grDevices, grid, gTable, isoband, MASS, mgcv, rlang, scales, stats, tibble, withr and dplyr. pheatmap” package was applied to construct heat maps and hierarchical clustering analyses. **Process and pathway enrichment analyses** were performed using METASCAPE^66^. Briefly, for each gene list, pathway and process enrichment analysis has been carried out with the following ontology sources: KEGG Pathway, GO Biological Processes, Reactome Gene Sets, Canonical Pathways, WikiPathways and PANTHER Pathway. All genes in the genome have been used as the enrichment background. Terms with a p-value < 0.01, a minimum count of 3, and an enrichment factor > 1.5 (the enrichment factor is the ratio between the observed counts and the counts expected by chance) are collected and grouped into clusters based on their membership similarities. More specifically, p-values are calculated based on the accumulative hypergeometric distribution, and q-values are calculated using the Benjamini-Hochberg procedure to account for multiple testings. **Protein Protein Interaction Analysis** was assisted by METASCAPE^66^ and CYTOSCAPE^67^. For each target gene list, protein-protein interaction enrichment analysis has been carried out with the following databases: STRING, BioGrid, OmniPath, InWeb_IM. Only physical interactions in STRING (physical score > 0.132) and BioGrid are used. The resultant network contains the subset of proteins that form physical interactions with at least one other member in the list. If the network contains more than 5 proteins, the Molecular Complex Detection (MCODE) algorithm has been applied to identify densely connected network components. Pathway and process enrichment analysis has been applied to each MCODE component independently, and the best-scoring term by p-value have been retained as the functional description of the corresponding component. **Venn diagrams** were performed using Bioinformatics and evolutionary genomics webtool (http://bioinformatics.psb.ugent.be/webtools/Venn/). **All Figs and cartoons** were edited and laid out using the open-source Vector graphics editor Inkscape (https://inkscape.org/)

