## Supplemental Figures for "The eukaryotic translation initiation factor eIF4E reprogrammes the splicing machinery and drives alternative splicing"

**Supp Fig 1. Comparison of eIF4E levels and localization in U2OS and NOMO cell line with primary specimens**

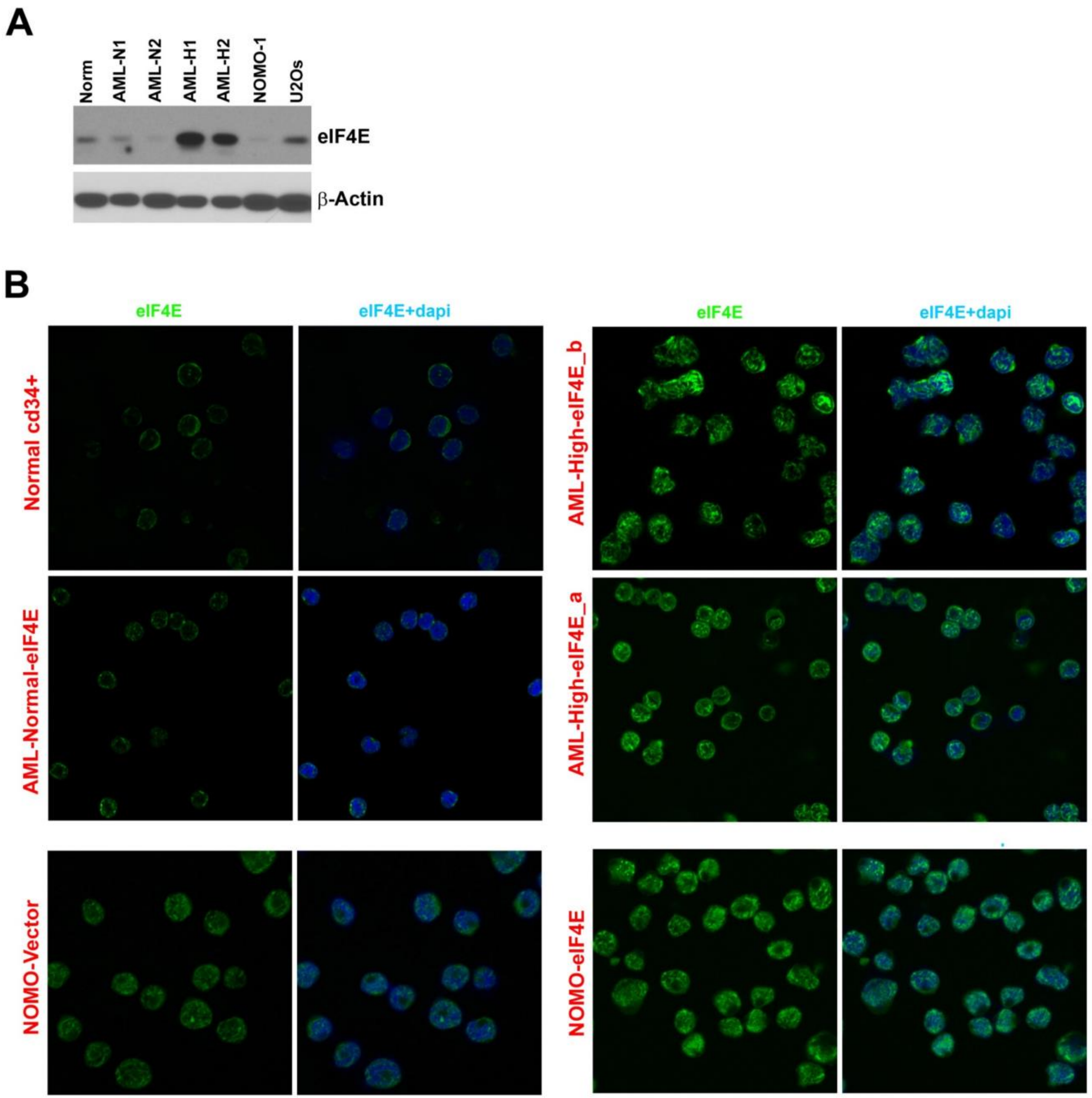

**C**

>NOMO1SF3B1_SF3B1Fw5

NNNNNNNNNGNTGCTGGTCTGGCTACTATGATCTCTACCNTGAGACCTGATATAGATAACATGGATGAGTA
TGTCCGTAACACAACAGCTAGAGCTTTTGCTGTTGTAGCCTCTGCCCTGGGCATTCCTTCTTTATTGCCCTTC
TTAAAAGCTGTGTGCAAAAGCAAGAAGTCCTGGCAAGCGAGACACACTGGTATTAAGATTGTACAACAGA
TAGCTATTCTTATGGGCTGTGCCATCTTGCCACATCTTAGAAGTTTAGTTGAAATCATTGAACATGGTCTTGT
GGATGAGCAGCAGAAAGTTNNGACCATCAGTGCTTTGGCCATTGCTGCCTTGGCTGAAGCAGCAACTCCT
TATGGTATCGAATCTTTTGATTCTGTGTTAAAGCCTTTATGGAAGGGTATCCGCCA

Equal mix of A and C (CGG=wt)

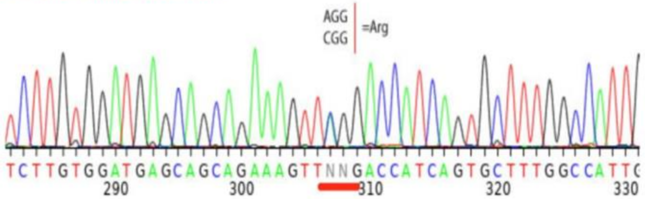


>NOMO1SF3B1_SF3B1Fw1

NNNNNNNNNNCTGNGTGCAANGCAAGAAGTCCTGGCAAGCGAGACACACTGGTATTAAGA
TTGTACAACAGATAGCTATTCTTATGGGCTGTGCCATCTTGCCACATCTTAGAAGTTTAG
TTGAAATCATTGAACATGGTCTTGTGGATGAGCAGCAGAAAGTTNGGACCATCAGTGCTT
TGGCCATTGCTGCCTTGGCTGAAGCAGCAACTCCTTATGGTATCGAATCTTTTGATTCTG
TGTTAAAGCCTTTATGGAAGGGTATCCGCCA

Equal mix of A and C (CGG=wt)

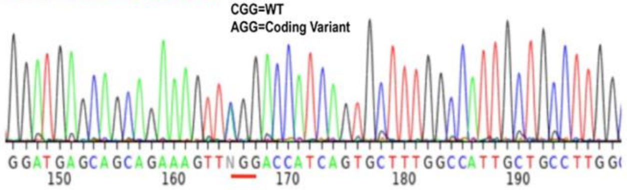


**rs16865307**

chr2:197402104 (GRCh38.p12); Gene ConsequenceSF3B1 : Missense Variant

Alleles:G>A / G>T

| Molecule type | Change | Amino acid[Codon] | SO Term |
|---|---|---|---|
| SF3B1 transcript variant 1 | NM_012433.4:c.2104C>T | R [CGG] > W [TGG] | Coding Sequence Variant |
| SF3B1 transcript variant 1 | NM_012433.4:c.2104C>A | R [CGG] > R [AGG] | Coding Sequence Variant |
| SF3B1 transcript variant 2 | NM_001005526.2:c. | N/A | Genic Downstream Transcript Variant |
| SF3B1 transcript variant 3 | NM_001308824.1:c. | N/A | Genic Downstream Transcript Variant |
| SF3B1 transcript variant X1 | XR_001738680.2:n.2149C>T | N/A | Non Coding Transcript Variant |
| SF3B1 transcript variant X1 | XR_001738680.2:n.2149C>A | N/A | Non Coding Transcript Variant |
| SF3B1 transcript variant X2 | XR_001738681.1:n. | N/A | Genic Downstream Transcript Variant |
| SF3B1 transcript variant X3 | XR_241302.2:n. | N/A | Genic Downstream Transcript Variant |
| splicing factor 3B subunit 1 isoform 1 | NP_036565.2:p.Arg702Trp | R (Arg) > W (Trp) | Missense Variant |
| splicing factor 3B subunit 1 isoform 1 | NP_036565.2:p.Arg702= | R (Arg) > R (Arg) | Synonymous Variant |

NOMO1-SRSF2_SRSFFw1

TATGGNTGCCNTGGACGGGGCCGTGCTGGACGGCCGCGAGCTGCGGGTGCAAATGGCGCGCTACGGCCGCCCCCCGGACTCACACCACA
GCCGCCGGGGACCGCCACCCCGCAGGTACGGGGGCGGTGGCTACGGACGCCGGAGCCGCAGCCCTAGGCGGCGTCGCCGCAGCCGATCC
CGGAGTCGGAGCCGTTCCAGGTCTCGCAGCA **Corresponds to WT**

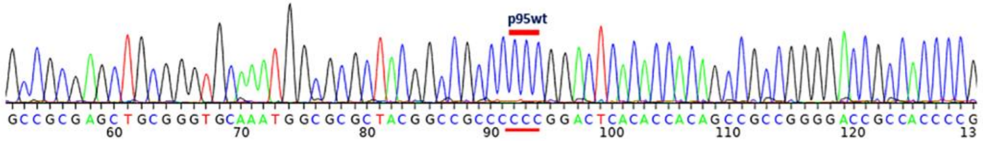


**NOMO1-U2AF35_U2AF35Fw4**

AAAGTCAACTGTTCATTTTATTTCAAAATTGGAGCATGTCGTCATGGAGACAGGTGCTCTCGGTTGCACAATAAACCGACGTTTAGCCAGAC
CATTGNCCTCTTGAACATTTACCGTAACCCTCAAAACTCTTCCCAGACTGCTGACGGNTTGCNCTGTGCCGTGAGCGATGTGGAGATGCAG

**Corresponds to WT**

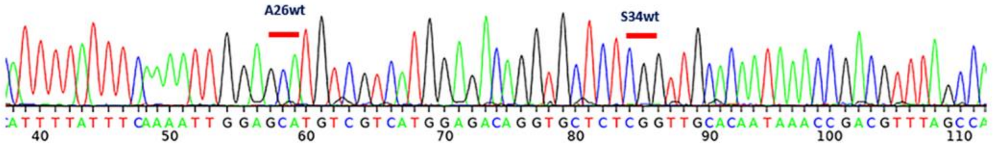


**NOMO_4EU2AF35_u2af35Q157R1**GCCGCAGCTCTCTGGAAATGGGCTTCAAATGCATGAAGTTGCAGAAGCCGCCTCGTGTGCATTC
TCCCATCTCATACTGACGGCAGCAGGCTTCTCTGAAGTCCGTCACGGGTGACAGCTCGGCGTGGATCGGCTGTCCATTAAACCAACGGTTA
TTCAAGTCAATCACAGCCTTTTCCGCATCTTCCTCACGGCGAAACTTGACGTACACGTTCCCCACCAGGTGGTCTCCCAGGTTGTCACAGA
CGTTCATCTCCTCTACTTCCCCATACTTCTCCTCCATTTCTGTAAAAACCTCCTCAAAAAACTCATCATAGTGTTCCTGCATCTCCACATCGCTC
ACGGCACAGCGCAAACCGTCAGCAGACTGGGAANAGNTTTGAGGGTTACGGTAAATGTTCAAGAGGGCAATGGTCTGGCTAAACGTCG
GTTTATTGTGCAACCGAGAGCACCTGTC TCCATGACGACATGCTCCAATTTTGAAATAAAATGAACAGNTGACTTTGTCTTTCTCGGT
GNCGAAGATGGANGNCANATACTCC

**Corresponds to WT**

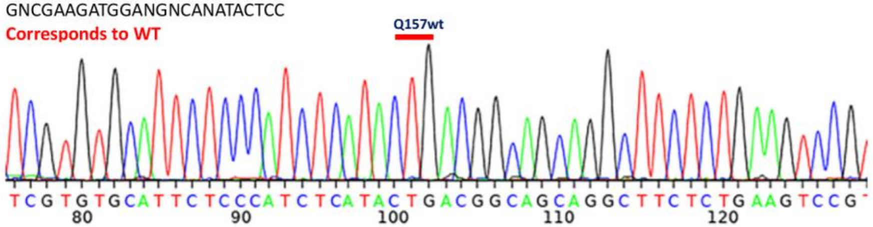

**Supp Fig 1.A.** WB analysis of endogenous eIF4E levels in U2OS and NOMO-1 cell lines compared to primary AML samples with high (AML-H) and normal (AML-N) eIF4E levels, as well as bone marrow mononuclear cells from healthy volunteers (Norm). β-Actin was used as a loading control. **B.** Localization of eIF4E in NOMO-1 Vector and eIF4E cells compared to primary AML specimens with high (AML-H) and normal (AML-N) eIF4E levels, as well as bone marrow mononuclear cells from healthy volunteers (Norm). Confocal micrographs of cells stained with anti-eIF4E antibodies (in green) and DAPI (in blue) as a nuclear marker. Single (eIF4E) and overlaid (eIF4E+DAPI) channels are shown. Micrographs are single sections through the plane of the cells with 63x magnification. Confocal settings were identical for all samples and thus the differences in staining intensity is related to the levels of eIF4E. **C.** Sequencing of SF3B1, SRSF2 and U2AF1 genes for NOMO-1 cells showing WT phenotype. SF3B1 showed heterozygous status with allele variant already reported. This allele does not alter the amino acids in the protein produced i.e. results in a silent mutation.

**Supp Fig 2. eIF4E regulates expression of SF factors and interacts with UsnRNAs**

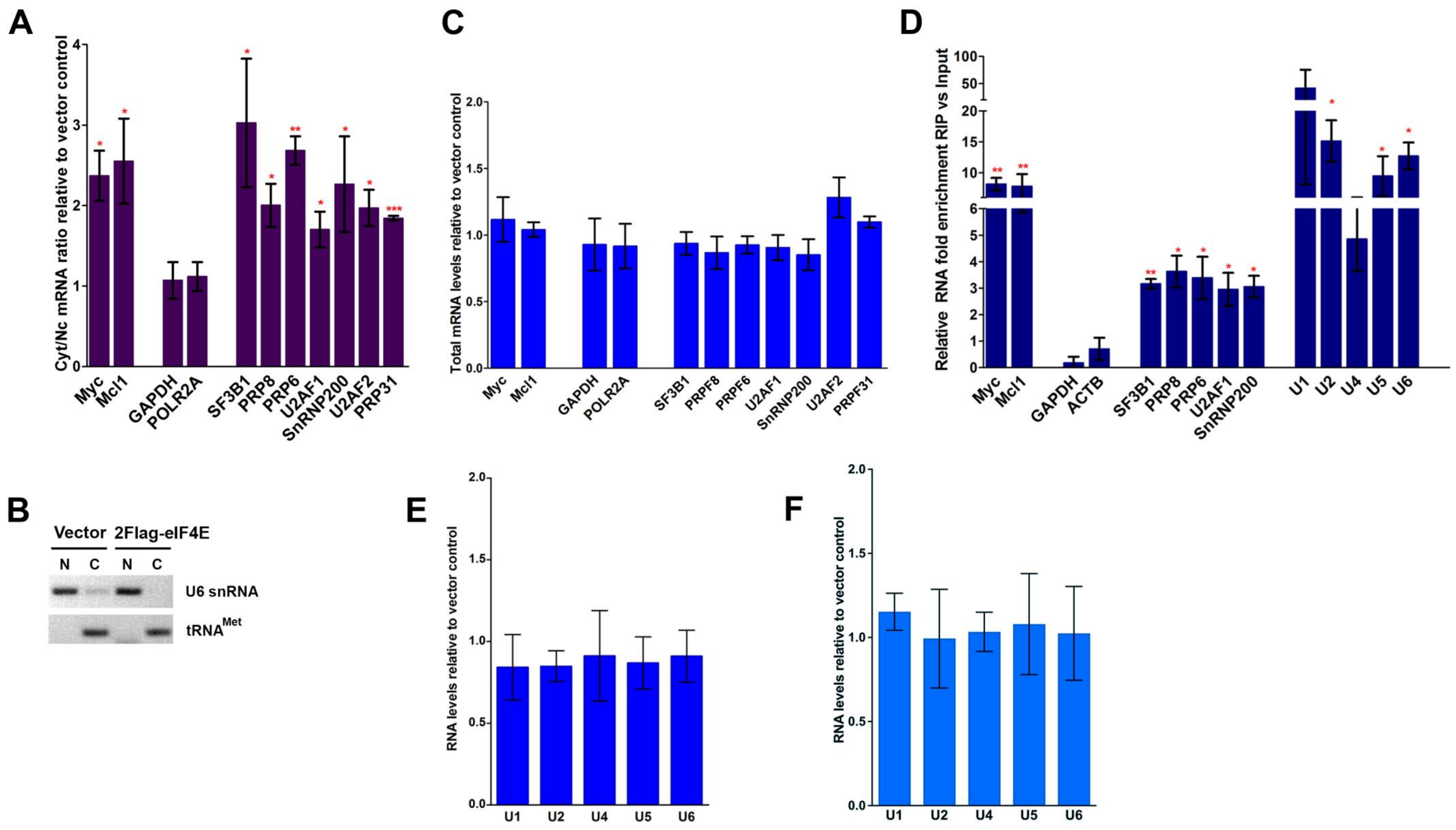

**Supp Fig 2. A.** RNA export assays for 2FLAG-eIF4E and Vector control U2OS cell lines. Transcript levels of targets in nuclear and cytoplasmic compartments were assessed by RT-qPCR with fractionation controls shown in Supp Fig 2B. Data were normalized to vector control and shown as a fold change. ACTNB was used as a loading control. The mean, standard deviation, and p-values were derived from three independent replicates i.e. form three different cell lines for each condition, with each carried out in triplicate (* $p<0.05$, ** $p<0.01$, *** $p<0.001$). **B.** Semi-qPCR for U6 snRNA and $tRNA^{Met}$ as controls for the cytoplasmic and nuclear fractions, respectively, corresponding to the export assay shown in A; N=nuclear fraction, C=cytoplasmic fraction. Representative experiment (out of 3 biological replicates) is shown. **C.** Total mRNA levels monitored by RT-qPCR corresponding to mRNA export assays shown in A. Data were normalized to vector control to calculate fold change. The mean and standard deviation, as well as p-values, were derived from three biological experiments, each carried out in triplicate. **D.** The enrichment of mRNAs and UsnRNAs in RIPs of endogenous eIF4E versus input RNAs from the nuclear fractions of NOMO-1 cells monitored by RT-qPCR. Data were normalized to input samples and presented as a fold change. The mean, standard deviation and p-values were derived from three independent experiments (each carried out in triplicate). Myc, and Mcl1 are known eIF4E nuclear targets and served as positive controls, while ACTB and GAPDH were used as negative controls. **E.** Total levels of UsnRNA in U2OS Vector and 2FLAG-eIF4E cells monitored by RT-qPCR. Data were normalized to vector control and shown as a fold change. The mean and standard deviation, as well as p-values were derived from three biological replicates each carried out in triplicate. **F.** Total levels of UsnRNA in NOMO-1 Vector and eIF4E cells monitored by RT-qPCR. Data were normalized to vector control and shown as a fold change. The mean and standard deviation, as well as p-values were derived from three biological replicates each carried out in triplicate.

**Supp Fig 3. Alternative Splicing reprogramming upon eIF4E overexpression is based solely on eIF4E levels and does not correlate to RNA levels in U2OS cells**

A

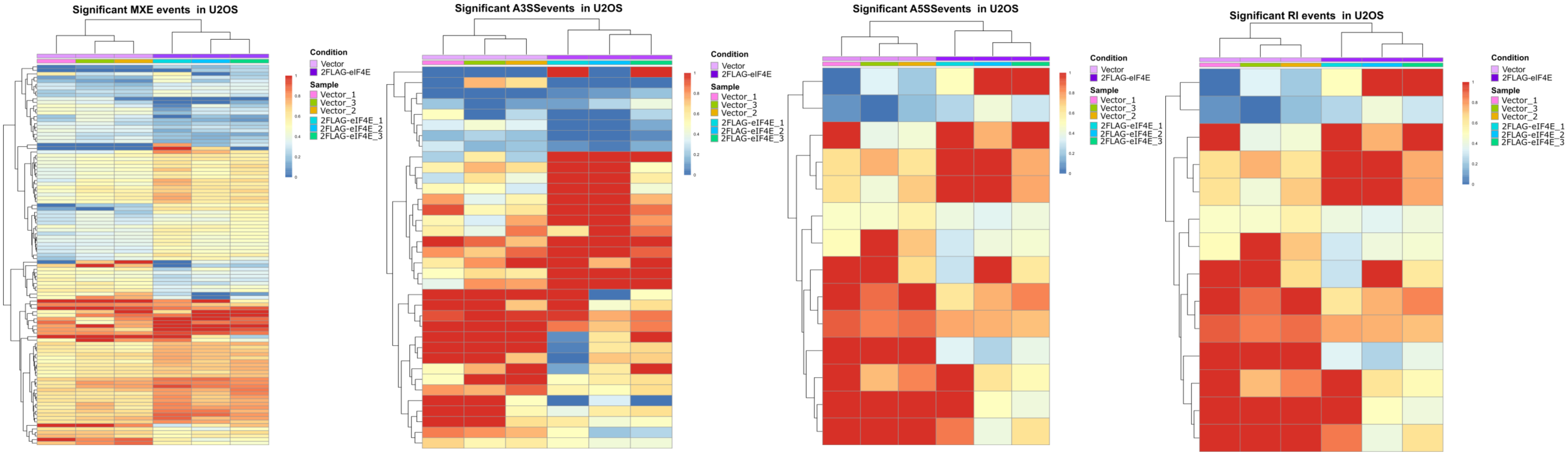

B

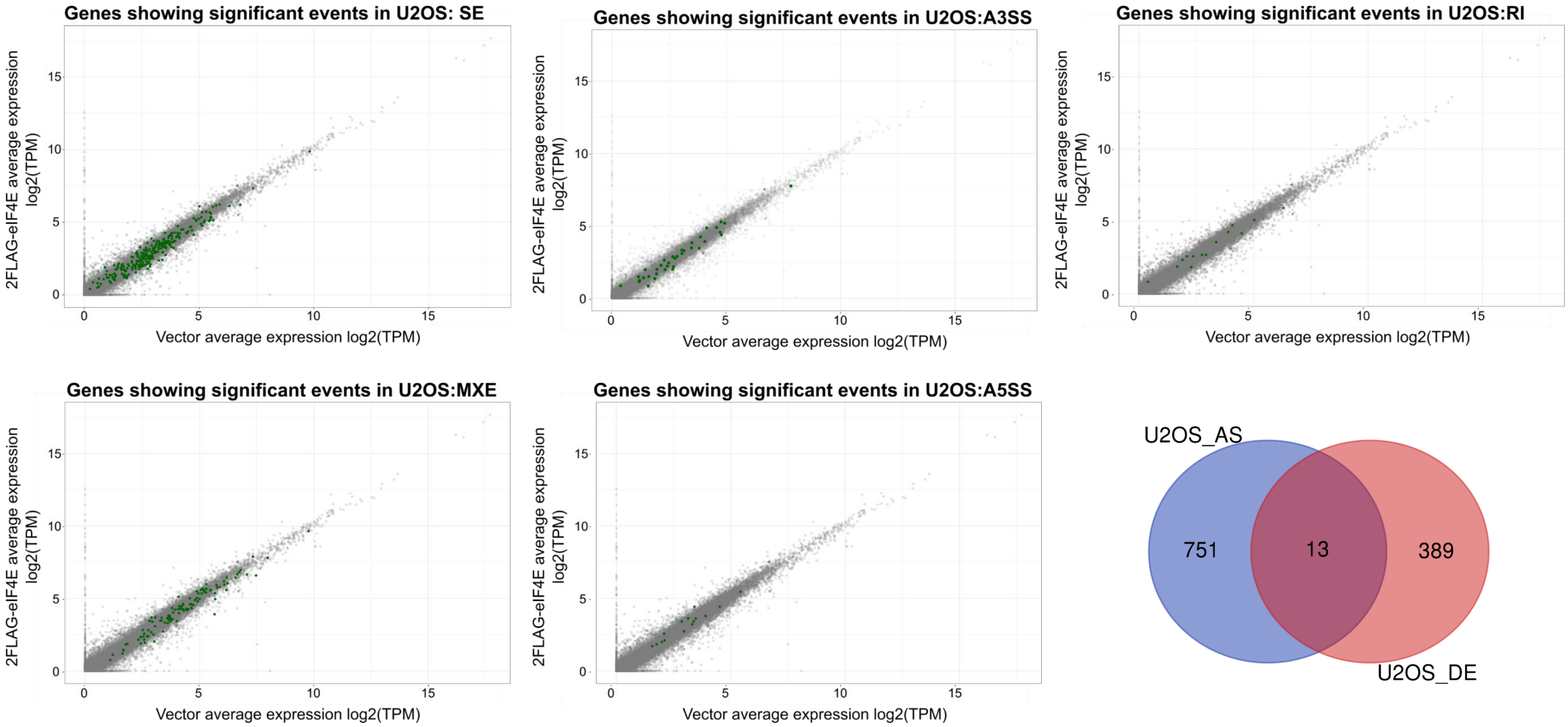

Genes showing significant events in U2OS: SE
Genes showing significant events in U2OS:A3SS
Genes showing significant events in U2OS:RI
Genes showing significant events in U2OS:MXE
Genes showing significant events in U2OS:A5SS
2FLAG-eIF4E average expression log2(TPM)
Vector average expression log2(TPM)
0
5
10
15
U2OS_AS
U2OS_DE
751
13
389

**Supp Fig 3. A.** Unsupervised hierarchal clustering indicates that events segregate solely on eIF4E levels in U2Os cells. The resulting heatmaps of inclusion levels for indicated event type are shown. Events considered have an FDR-adjusted p-value <0.1 and an absolute inclusion value >0.1. SE events are shown in Fig 3C. **B.** Alternatively spliced targets are not characterized by differential RNA expression in 2FLAG-eIF4E versus Vector U2OS cells: Scatter plots show average $\log_2$(TPM) for each condition. Targets that have one or more splicing events are highlighted in green. Lower right panel. Venn diagram showing the overlap between differentially expressed genes (DE) and alternatively spliced targets (AS) is low, with only 13 targets in common.

**Supp Fig 4. High eIF4E levels are correlated with low survival in AML patients and with alternative splicing reprogramming, independently of differential gene expression**

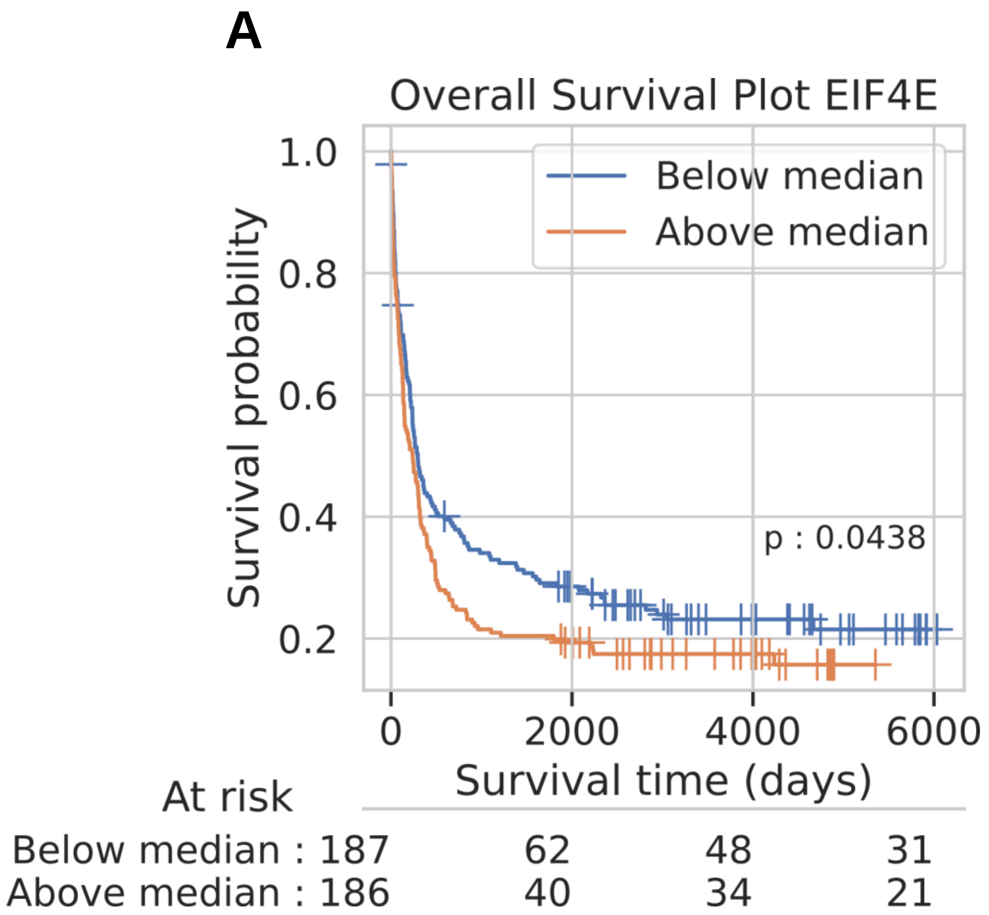


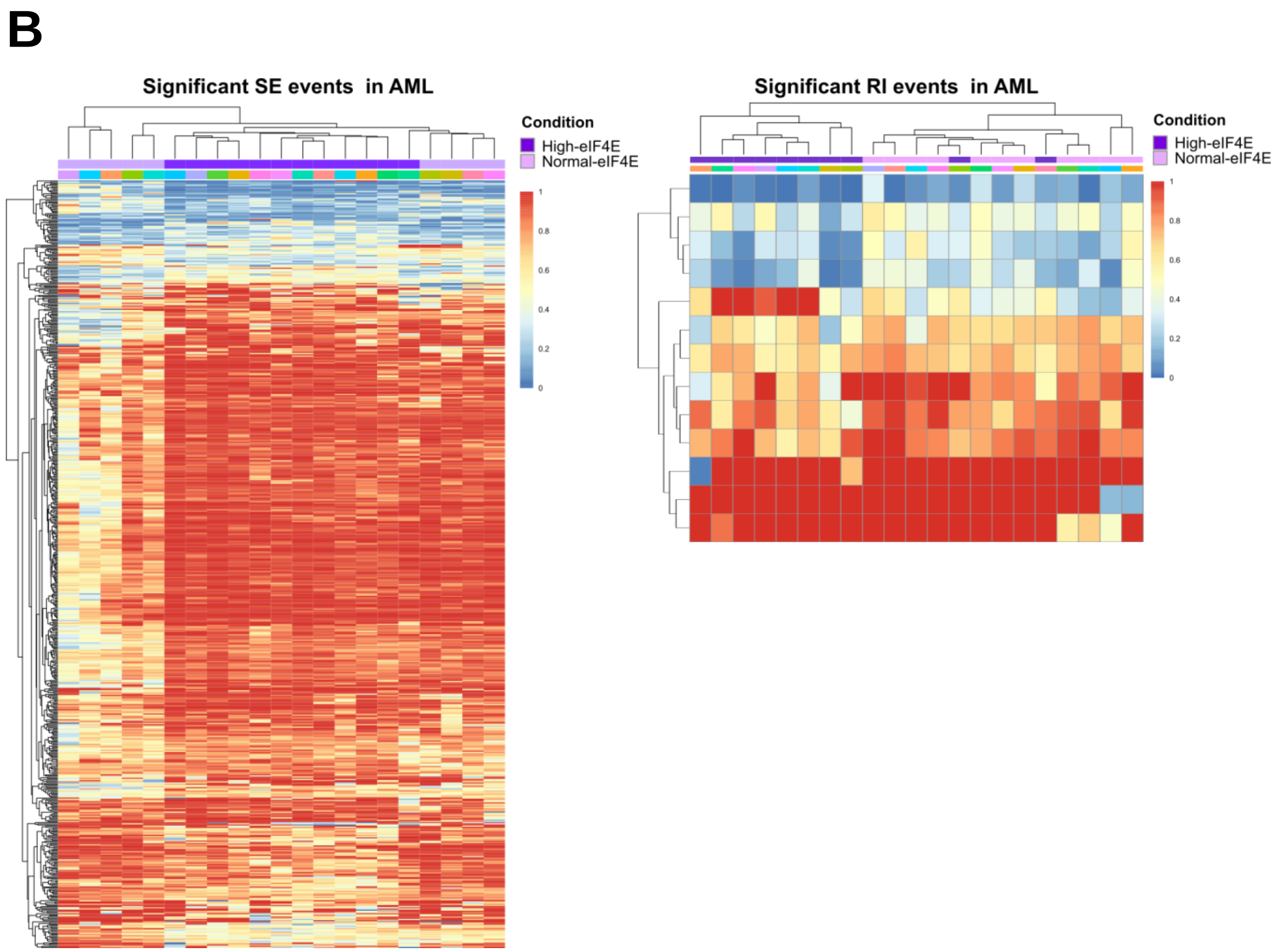

C

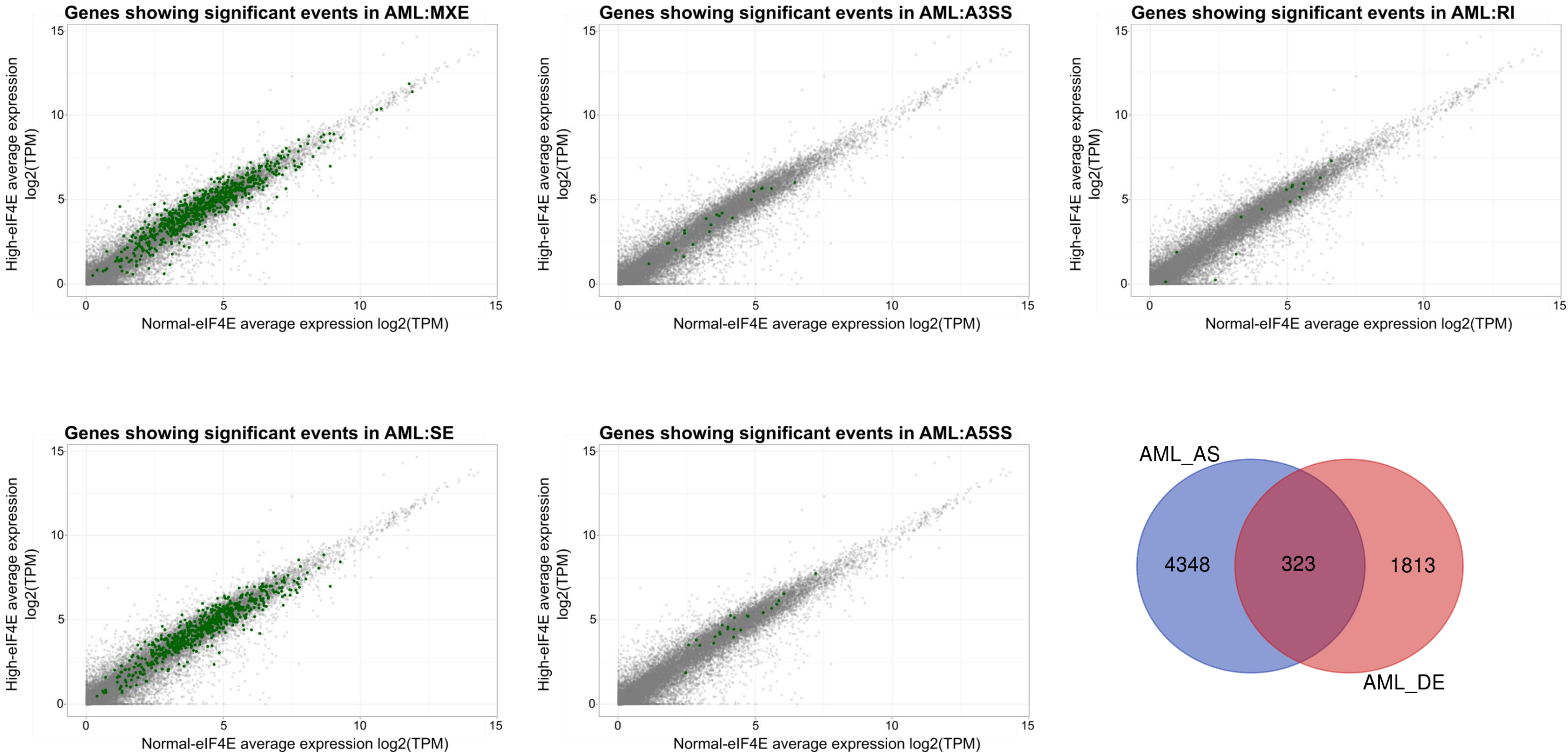
Genes showing significant events in AML:MXE
Genes showing significant events in AML:A3SS
Genes showing significant events in AML:RI
Genes showing significant events in AML:SE
Genes showing significant events in AML:A5SS
High-eIF4E average expression log2(TPM)
Normal-eIF4E average expression log2(TPM)
0
5
10
15
AML_AS
AML_DE
4348
323
1813

**Supp Fig 4. A.** Survival analysis of the Leucegene cohort. The cohort was divided in two groups based on eIF4E expression levels (below the median, blue and above the median, orange). Log Rank test was performed and shows a significant difference in survival between the two groups (p-value<0.05). **B.** Unsupervised hierarchal clustering of indicated AS events in primary AML specimens. The resulting heatmaps show that events segregate on eIF4E levels (High-eIF4E versus Normal-eIF4E specimens), but in AML there is more than eIF4E group; whereas in RI events, clustering on eIF4E levels is evident but there are two outliers that behave more like Normal-eIF4E AML. Heatmaps of inclusion levels for SE and RI events are shown here and for MXE events in Fig4E. Events considered have an FDR-adjusted p-value <0.1 and an absolute inclusion value >0.1. **C.** Alternatively spliced targets are not differentially expressed in High-eIF4E versus Normal-eIF4E AML patient' cells: Scatter plots showing average log2 (TPM) for each condition. Targets that have one or more splicing event(s) are highlighted in green. Venn diagram showing the overlap between differentially expressed genes (DE) and alternatively spliced targets (AS) is low, with only 7% overlap.

**Supp Fig 5. Characterization of eIF4E-dependent alternative splicing events in U2OS cells.**

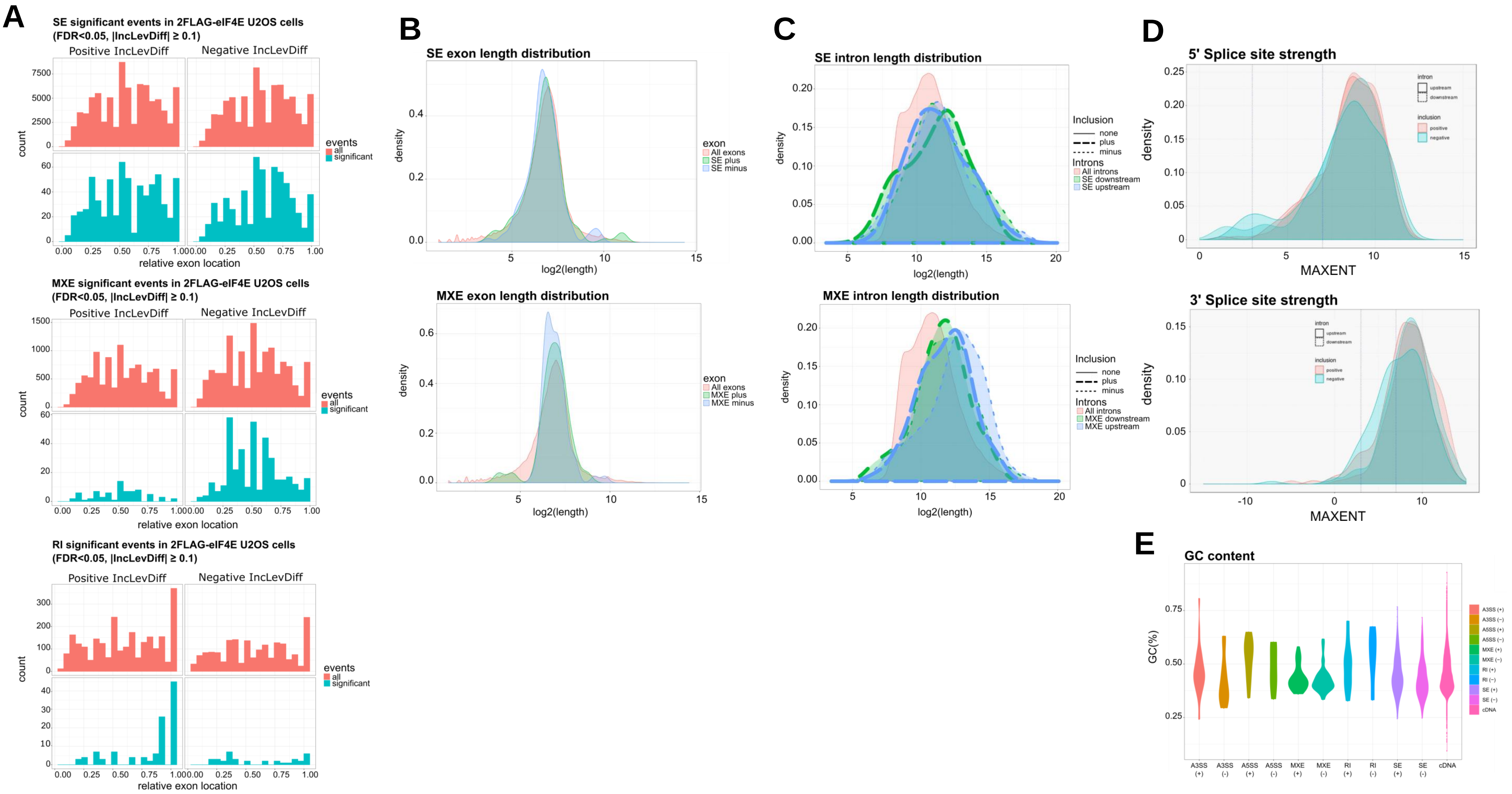

**Supp Fig 5. A.** Relative location of significant splicing events along the length of the transcripts with each transcript divided into equal fractions where 0.00 indicates the 5' end and 1.00 the 3' end. Events are shown for all transcripts (pink) and those that were significantly different between 2FLAG-eIF4E and Vector cells (cyan). To capture the most robust differences, FDR adjusted p-values<0.05 and absolute inclusion level differences >0.1 are shown, and data were segregated based on the sign of the inclusion levels differences "Positive IncLevDiff" and "Negative IncLevDiff". **B.** The length distribution of exons involved in alternate splicing events (FDR<0.1, absolute inclusion level difference > 0.1) compared to all exons. Plus refers to positive inclusion level differences (green), negative differences (blue) and for comparison, distribution of lengths for all exons (orange). **C.** The length distribution of introns involved in alternate splicing events (FDR<0.1, absolute inclusion level differences > 0.1) compared to length distribution for all introns (in orange). The intron is downstream (green) or upstream (blue) of the splicing event. Plus refers to positive inclusion level differences and minus refers to negative differences. **D.** Prediction of splice site strength based on sequence analysis around splice site events using MAXENT. Positive refers to positive inclusion level differences (orange), minus refers to negative inclusion level differences (blue). The position of the 5' splice site upstream (solid line) or downstream (dashed line) of the event is shown. **E.** Comparison of GC content in introns involved in significant events (FDR-adjusted p-value <0.1, absolute inclusion level difference > 0.1) compared with all cDNA. The GC content for cDNA was computed on all transcripts from Gencode version 32 of the human genome. "+" indicates positive inclusion level difference, and "–" a negative one.

**Supp Fig 6. Characterization of eIF4E-dependent alternative splicing events in AML specimens.**

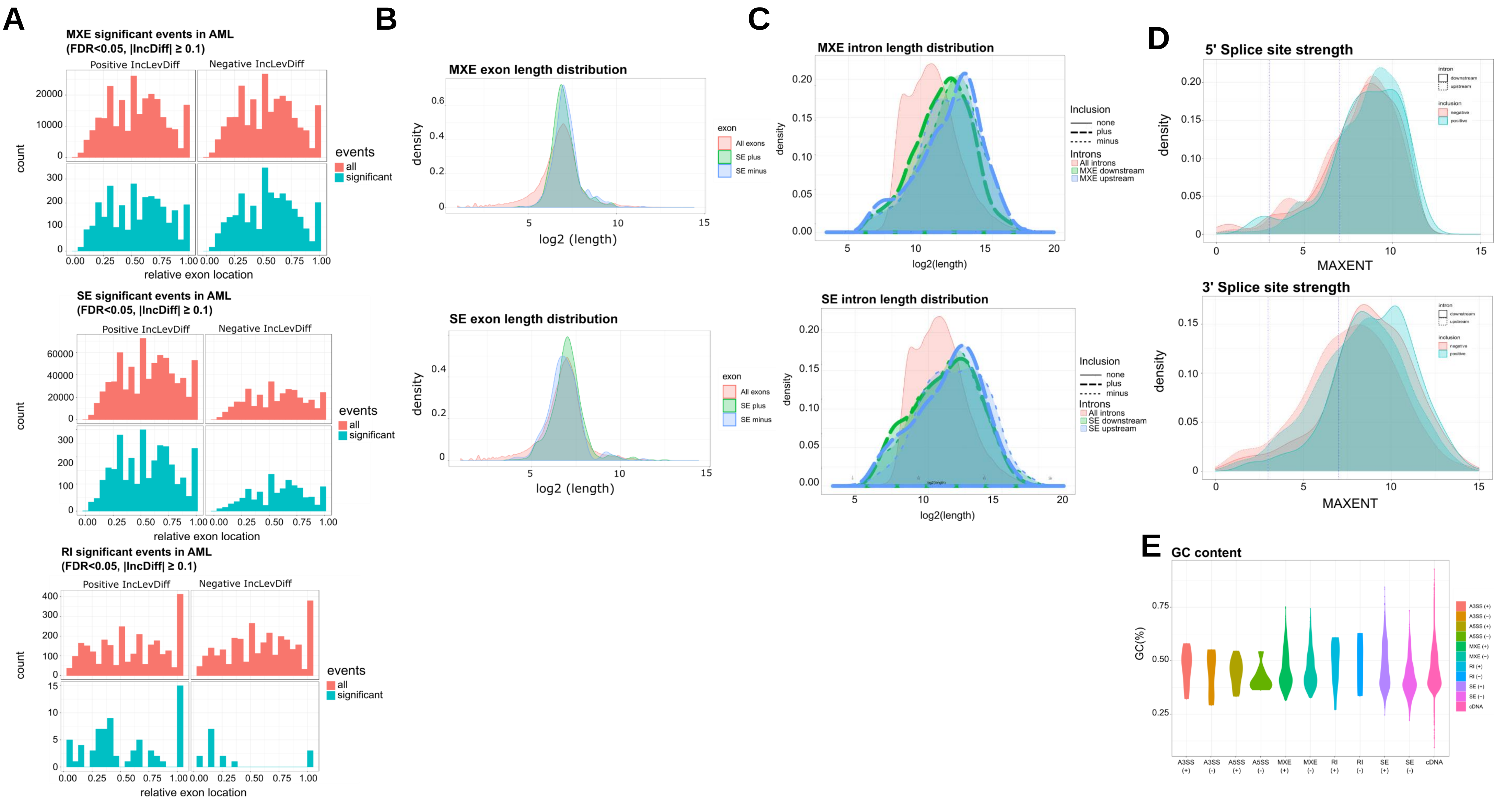

**Supp Fig 6. A.** Relative location of significant splicing events along the length of the transcripts with each transcript divided into equal fractions where 0.00 indicates the 5' end and 1.00 the 3' end. Events are shown for all transcripts (pink) and those that were significantly different between High-eIF4E and Normal-eIF4E cells (cyan). To capture the most robust differences, FDR adjusted p-values<0.05 and absolute inclusion level differences >0.1 are shown, and data were segregated based on the sign of the inclusion levels differences "Positive IncLevDiff " and "Negative IncLevDiff ". **B.** The length distribution of exons involved in alternate splicing events (FDR<0.1, absolute inclusion level difference > 0.1) compared to all exons. **C.** The length distribution of introns involved in alternate splicing events (FDR-adjusted p-value<0.1, absolute inclusion level differences > 0.1) compared to length distribution for all introns (in orange). The intron is downstream (green) or upstream (blue) of the splicing event. Plus refers to positive inclusion level differences and minus refers to negative differences. **D.** Prediction of splice site strength based on sequence analysis around splice site events using MAXENT. "positive" refers to positive inclusion level differences (orange), "negative" refers to negative differences (blue). The position of the 5' splice site upstream (solid line) or downstream (dashed line) of the event is shown. **E.** Comparison of GC content in introns involved in significant events (FDR-adjusted p-value <0.1, absolute inclusion level difference > 0.1) compared with all cDNA. The GC content for cDNA was computed on all transcripts from Gencode version 32 of the human genome. "+" indicates positive inclusion level difference, and "–"a negative one.

**Supp Fig 7. Comparisons of U2Os and AML related rMATS analyses with published datasets.**

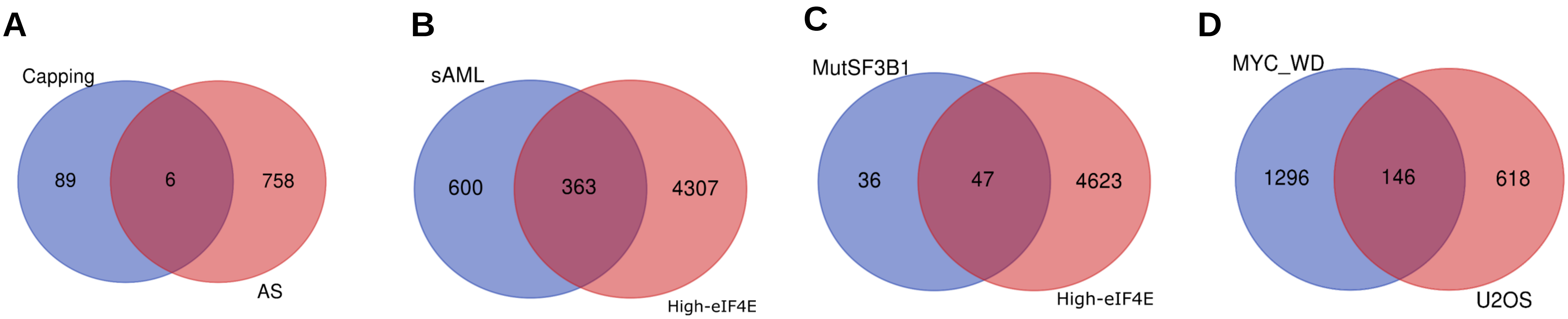


**Supp Fig 7. A.** Venn diagram showing the overlap between eIF4E-Capping[7] and eIF4E-AS targets identified here. **B.** Comparison of alternatively spliced targets in secondary AML (sAML) with AS transcripts in High-eIF4E AML patients' samples here. **C.** Venn diagram showing the overlap between alternatively spliced targets in AML samples harboring SF3B1 mutations (MutSF3B1, mainly the hotspot mutation K700)[26] and AS targets in High-eIF4E AML patient's samples. **D.** Comparison of splicing targets following Myc withdrawal in Myc/myrAKT1 Prostate cancer cells (MYC-WD)[31] with alternatively spliced targets upon eIF4E overexpression.

**Supp Fig 8. eIF4E positioned to influence splicing through physical interactions with splicing components and substrate RNAs (direct model), and/or by regulation of SFs levels (indirect model).**

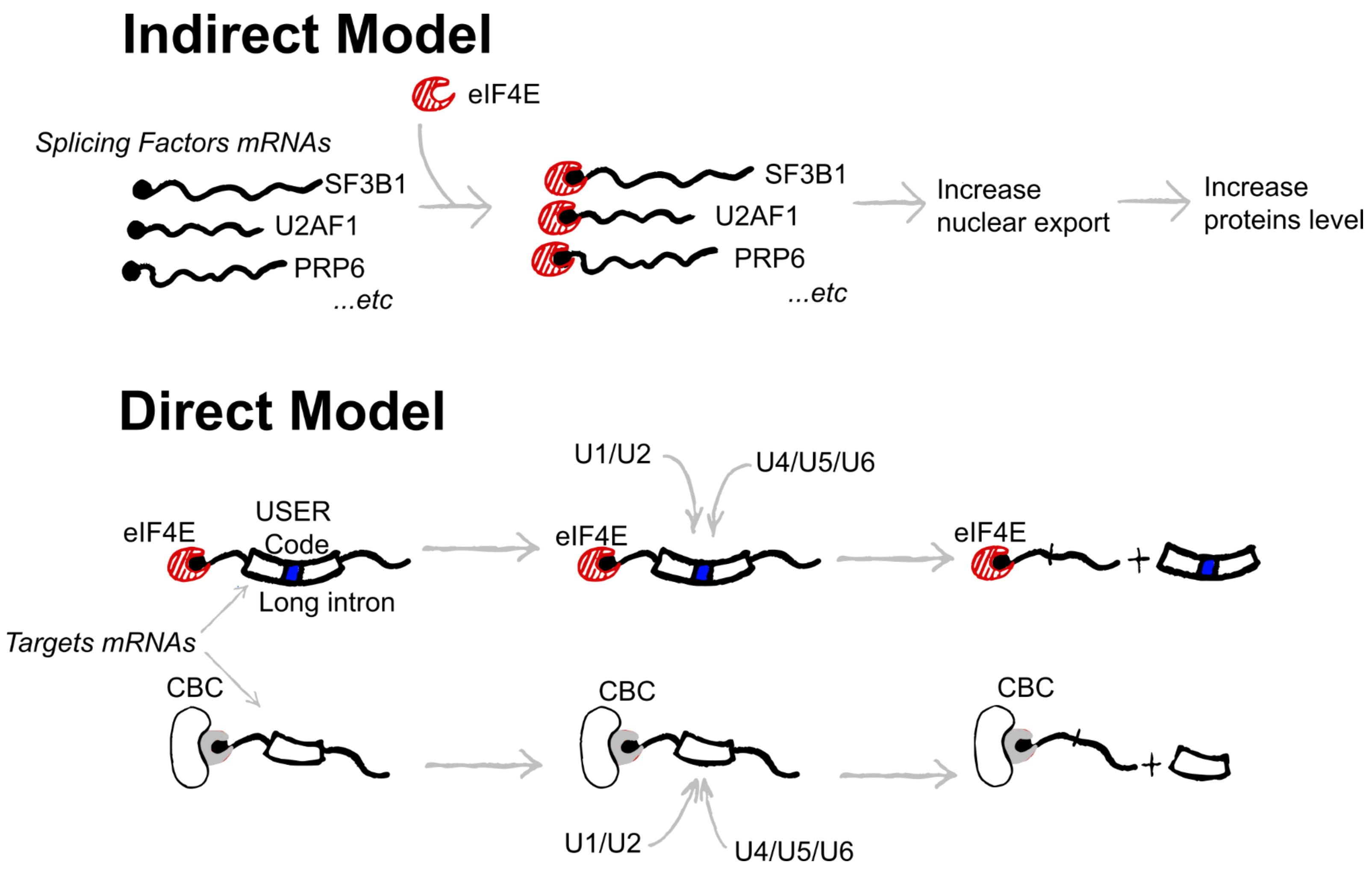
