## Supplementary material for "The eukaryotic translation initiation factor eIF4E reprogrammes the splicing machinery and drives alternative splicing": Supp Table 14

| Cell type- Event | Upstream + | p-val vs avg | Downstream + | p-val vs avg | Upstream - | p-val vs avg | Downstream - | p-val vs avg | Average expressed introns (gene TPM>=1) |
|---|---|---|---|---|---|---|---|---|---|
| U2OS- SE | 9670 | 2.12E-03 | 7225 | 1.67E-05 | 11477 | 5.62E-04 | 11153 | 2.04E-04 | 5047 |
| U2OS- MXE | 7421 | 9.96E-05 | 6069 | 1.57E-03 | 13741 | < 2.2E-16 | 9025 | 3.79E-09 | 5047 |
| AML- SE | 6962 | < 2.2E-16 | 6334 | 6.33E-15 | 9759 | 7.3E-13 | 7481 | 1.27E-6 | 4634 |
| AML- MXE | 9870 | < 2.2E-16 | 6845 | < 2.2E-16 | 9955 | < 2.2E-16 | 9415 | < 2.2E-16 | 4634 |

**Supp Table 14. eIF4E sensitive splicing events are generally associated with longer introns.** Tabulation of results observed in Supp Fig 5C and Supp Fig 6C. The predominant splicing events for U2Os and AML were MXE and SE and thus were focussed on here. Upstream and downstream introns were evaluated based on the sign of the inclusion level difference between 2FLAG-eIF4E and Vector cells relative to Normal-eIF4E and High-eIF4E AML specimens. For comparison, average intron length (avg) in U2Os or AML cells are shown. Lengths are given in base pairs. p-values were computed using Kolmogorov–Smirnov test to determine if the two underlying distributions differ significantly.
