## Supplementary material for "The eukaryotic translation initiation factor eIF4E reprogrammes the splicing machinery and drives alternative splicing": Supp Table 21

| Normal-eIF4E samples | High-eIF4E samples |
|---|---|
| 02H003 | 04H118 |
| 02H017 | 05H008 |
| 05H078 | 06H135 |
| 05H128 | 10H053 |
| 06H020 | 10H115 |
| 07H045 | 11H014 |
| 08H042 | 11H145 |
| 09H013 | 11H183 |
| 11H107 | 12H173 |
| 12H030 | 13H056 |
| 13H053 | |

**Supp Table 19.** List of AML samples (Normal-eIF4E and High-eIF4E) used in these studies from Leucegene.ca. None of these specimens had known mutations associated with AML in SF3B1, U2Af1 and SRSF2 as described in the text.
