## Supplementary material for "The eukaryotic translation initiation factor eIF4E reprogrammes the splicing machinery and drives alternative splicing": Supp Table 22

**Supp Table 22. List of primers**

| |
|---|
| **ActinBF:** GCATGGAGTCCTGTGGCATCCACG |
| **ActinBR:** GGTGTAACGCAACTAAGTCATAG |
| **Mcl1Fw:** GGCAGTCGCTGGAGATTAT |
| **Mcl1 Rv:** CAACCCGTCGTAAGGTCTC |
| **MycFw**: CTTCTCTGAAAGGCTCTCCTTG |
| **MycRv**: GTCGAGGTCATAGTTCCTGTTG |
| **POL2ARFw:** TGACTGCCAACACAGCCATCTACT |
| **POL2ARRv:** GGGCCACATCAAAGTCAGGCATTT |
| **GAPDHFw:** GAAGGTGAAGGTCGGAGTC |
| **GAPDHRv:** GAAGATGGTGATGGGATTTC |
| **18S Fw:** CGG CGA CGA CCC ATT CGA AC |
| **18S Rv:** GAA TCG AAC CCT GAT TCC CCG TC |
| **U6Fw:** CGCTTCGGCAGCACATATAC |
| **U6Rv:** AAAATATGGAACGCTTCACGA |
| **tRNAMetFw:** 5′-AGC AGA GTG GCG CAG CGG-3′, |
| **tRNAMetRv:** 5′-GAT CCA TCG ACC TCT GGG TTA-3′; |
| **FAAP20CommFw:** ATGCGGTGGTTAAGGATACAG |
| **FAAP20CommRv:** CAGAGTTCGAAGGCTTTGAGTA |
| **FAAP20SplicFw:** TCACTCAGAGGCATGAGGT |
| **FAAP20SplicRv:** CAGAGCTGAGGCTTCCTAGA |
| **IKBKBCommFw:** CAGAATCATCCATCGGGATCTAA |
| **IKBKBCommRv:** ATCCAGCTCCTTGGCATATC |
| **IKBKBSplicFw:** GGGCCTGGGAAATGAAAGA |
| **IKBKBSplicRv:** GGTGGGTCAGCCTGATTG |
| **MAPK8IP3CommFw:** GACATGTGTCCAGAGACCCG |
| **MAPK8IP3CommRv:** GGTACAGGGAGTCGGTGTTG |
| **MAPK8IP3SplicFw:** GTGCGCGATGATTTCTTTGG |
| **MAPK8IP3SplicRv:** AGGTCATTCTTCACCACATTCA |
| **IL4RCommFw:** ACAGTTCACACCAATGTCTCC |
| **IL4RCommRv:** TGTTGACTGCATAGGTGAGATG |
| **IL4RCommFw:** AGC AGG GGC GCG CAG GTG CCT T |
| **IL4RCommRv:** AGG TCA CGT TAT AGA TTC TGAAA |
| **IL1bCommFw:** ATGGACAAGCTGAGGAAGATG |
| **IL1bCommRv:** CCCATGTGTCGAAGAAGATAGG |
| **IL1bSplicFw:** CAGCCAATCTTCATTGCTCAAG |
| **IL1bSplicRv:** AGGAGCACTTCATCTGTTTAGG |
| **PRP8Fw:** AACTGGTATCGGGAGCATTG |
| **PRP8Rv:** TCAGGGCATTCAGCACATAG |
| **PRP6Fw:** CCCTGTTCCTGACAGTTTCTT |
| **PRP6Rv:** CACCTGGATAGGGTGTGTTAAG |
| **SF3B1Fw:** GGTGGAAGTGACAGCAGATT |
| **SF3B1Rv:** CTGACCAAGCAAACTCGTAGAT |
| **SnRNP200Fw:** AAGAAGAGCCAAGCGAAGAA |

| |
|---|
| **SnRNP200Rv:** GATGCCCACCATCTCATCAA |
| **U2AF2Fw:** AGAGCTGCTGACATCCTTTG |
| **U2AF2Rv:** GATGTCCACGTACTCACAGAAG |
| **U2AF1Fw:** CGTCAGTATGAGATGGGAGAATG |
| **U2AF1Rv:** GGATCGGGATCTTGATCTATGC |
| **PRP31Fw:** CTGAGGCCAACCAGAAGTATT |
| **PRP31Rv:** GCAGTCATTCAGGTGGACATA |
| **U1snRNAFw:** GGAGATACCATGATCACGAAGG |
| **U1snRNARv:** CCACAAATTATGCAGTCGAGTTT |
| **U2snRNAFw:** CGTCCTCTATCCGAGGACAATA |
| **U2snRNARv:** GTACTGCAATACCAGGTCGATG |
| **U4snRNAFw:** TCGTAGCCAATGAGGTTTATCC |
| **U4snRNARv:** GCCAATGCCGACTATATTTCAAG |
| **U5snRNAFw:** CTGGTTTCTCTTCAGATCGCATAA |
| **U5snRNARv:** AGACTCAGAGTTGTTCCTCTCC |
| **IKBKBInr2Ex3Fw:** CAGGAGGTGATTGCAGGTAA |
| **IKBKBInr2Ex3Rv:** GCTCACCTGTTTCCTACAGAA |
| **IL4REx2Intr2Fw:** CATGCCTATAATCCCAGCACTT |
| **IL4REx2Intr2Rv:** GCCACCATGCCCAACTAAT |
| **MAPK8IP3Intr9Ex10Fw:** CGCTTCTCTTCTCTCCCTTTC |
| **MAPK8IP3Intr9Ex10Rv:** AGAAATCATCGCGCACTAGG |
| **FAAP20Intr6Fw:** AGCGCTGGCATTCTTGT |
| **FAAP20Intr6Rv:** CACCCTACACACTCAGGTTTC |
| **IL1BIntr1Fw:** AAC TAG GTG CTA AGG GAG TCT |
| **IL1BIntr1Rv:** GAG AGG GAG AGA CAG AGA AAG A |
